# Functional harmonics reveal multi-dimensional basis functions underlying cortical organization

**DOI:** 10.1101/699678

**Authors:** Katharina Glomb, Morten L. Kringelbach, Gustavo Deco, Patric Hagmann, Joel Pearson, Selen Atasoy

## Abstract

The human brain consists of functionally specialized areas, which flexibly interact and integrate forming a multitude of complex functional networks. However, the nature and governing principles of these specialized areas remain controversial: a distinct modular architecture versus a smooth continuum across the whole cortex. Here, we demonstrate a candidate governing principle ubiquitous in nature, that resolves this controversy for the brain at rest, during perception, cognition and action: functional harmonic modes. We calculated the harmonic modes of the brain’s functional connectivity, called *“functional harmonics”*, from functional magnetic resonance imaging (fMRI) data in resting state of 812 participants. Each functional harmonic provides an elementary pattern of brain activity with a different spatial frequency. The set of all functional harmonics - ordered according to their spatial frequencies - can reconstruct any pattern of brain activity. The activity patterns elicited by 7 different tasks from the Human Connectome Project can be reconstructed from a very small subset of functional harmonics, suggesting a novel relationship between task and resting state brain activity. Further, the isolines of the continuous functional harmonic patterns delineate the borders of specialized cortical areas as well as somatotopic and retinotopic organization. Our results demonstrate a candidate scalable governing principle for functional brain organization, resolving the controversy between modular versus gradiental views, and demonstrate that a universal principle in nature also underlies human brain cortical organization.

## Introduction

The human brain is topographically organized into functionally specialized brain areas^1^. Integration of these areas in various different constellations allows for the immense complexity of human brain function^2^. Despite remarkable progress in mapping the brain into functionally meaningful subdivisions, known as cortical areas^3, 4^, and in identifying functionally relevant combinations of these areas forming the functional networks of the brain^5^, the principles governing this functional segregation and integration in the human brain have remained unknown. Here we demonstrate that the brain activity underlying higher order human cognition can be decomposed into fundamental building blocks of *harmonic modes*, which are ubiquitous in nature. Our decomposition of task-based brain activity maps into the set of functional harmonics reveals that each task can be described by a very small subset of functional harmonics, suggesting that the functional harmonics are the fundamental building blocks of not only resting state activity, but crucially of the brain activity underlying cognitive functions controlling human behaviour. As an example, our results show how the brain activity related to movement of the right hand and trunk can be decomposed into separate harmonic modes that share the somatosensory (functional harmonic 7) but involve either the hand (functional harmonic 11) or the trunk (functional harmonic 3), respectively.

The topographic organization of the brain into functionally specialized areas is one of its fundamental properties, observed in evolution as early as the last common ancestor of vertebrates^4, 6^. The individuality of each brain area is determined by its functional specification, its microstructure (cyto- and myeloarchitecture)^4^, and its inter- and intra-area connectivity^3^. Significant effort in neuroscience has been directed towards subdividing the brain into adjoining parcels, which are assumed to have uniform functional specification and homogeneous connectivity^3, 4^. A multitude of functionally distinct brain areas coordinate through synchronous fluctuations in their activity^7^. Coherent oscillations among distinct brain areas have been shown to be another evolutionarily conserved aspect of brain activity^8^. The overlap of the networks formed through these spontaneous system oscillations, termed the functional connectivity patterns, with the functional networks of the human brain identified by various sensory, motor, and cognitive task paradigms^9–12^, strongly indicates their relevance for the brain’s functionality, as well as behavior.

However, this modular view of brain organization, where separate, adjoining brain areas with uniform functionality and homogeneous structural connectivity integrate into functional networks through coherent oscillations, has been challenged by the presence of gradually varying boundaries between brain areas suggesting a degree of transition instead of sharply separated brain areas^13^, as well as by the existence of topographic mappings, which characterize the differences within a functionally specific brain area^14–16^. Topographic mappings including retinotopy^14^, somatotopy^15^, tonotopy^16^, show that representation of our visual field, body and auditory frequency spectrum are spatially continuously represented across the areas of the primary visual, somatomotor and auditory cortices, respectively, challenging the assumption of uniform functionality within the determined brain areas and demonstrating a smoothly varying functionality^13^.

As an alternative, theoretical work^17, 18^ and recent experimental findings^13^ suggested a “gradiental perspective”, where the functional organization of the cortex is argued to be continuous, interactive and emergent as opposed to mosaic, modular and prededicated^17^. Similar to the smoothly varying functionality of primary sensory and motor areas, association cortices functioning as integration centres for more complex or elaborated mental processes are hypothesized to emerge from the convergence of information across sensory modalities^18^ with increasing spatial distance on the cortex from the highly functionally specialized primary cortices^19^. Supporting this hypothesis, a principal connectivity gradient of cortical organization in the human connectome has been identified, where the functional networks of the human brain are located according to a functional spectrum from perception and action to more abstract cognitive functions^13^. Although converging evidence^13, 20, 21^ supports the continuous and emergent view of cortical organization, the principles underlying the functional organization in the brain remain largely unknown.

Here we demonstrate how the modular functional architecture emerges from the continuous gradiental organization of the human cortex, thus resolving a long-standing controversy in neuroscience. In particular, we show that the human brain’s functional organization across various scales is governed by the natural principle of harmonic modes, where the isolines of these harmonic modes capture specialized functional areas. Remarkably, the principle of harmonic modes underlies a multitude of physical and biological phenomena including the emergence of harmonic waves (modes) encountered in acoustics^22^, optics^23^, electron orbits^24, 25^, electro-magnetism^26, 27^ and morphogenesis^28, 29^. The principle of harmonicity is also respected in the human brain across multiple scales, ranging from the ocular dominance patterns of the early visual areas^30^, visual hallucinations^31–34^, to the organization of cortical and thalamic tissues^35, 36^. On the macroscopic scale, harmonic modes of a circle^37^ and of a sphere have been proposed to underlie cortical communication observed in electroencephalogram (EEG)^37^ and in functional magnetic brain imaging (fMRI)^38^. Moreover, harmonic modes of the structural connectivity of the human brain, i.e. Laplace eigenfunctions on the human connectome, have been found to predict disease propagation in dementia^39^, as well as the collective dynamics of cortical activity at the macroscopic scale revealing the resting state networks^36^.

In this work, we uncover the spatial shapes of the harmonic modes emerging from synchronous hemodynamic fluctuations in large scale brain activity as measured in fMRI, by solving the time-independent (standing) wave equation^36, 40^ on the functional connectivity (FC) of the human brain. These harmonic modes, called *“functional harmonics”*, decompose the communication structure (i.e. FC) of the human brain into a hierarchical set of (spatial) frequency-specific communication channels, which naturally emerge from coherent, spontaneous brain activity. The functional harmonics unveil both, the principal connectivity gradient^13^, as well as cortical parcellations^3^. Overall, our results show that the functional segregation and integration in the brain, enabling complex human behaviour, are governed by this multi-dimensional harmonic representation.

## Results

### Estimation of functional harmonics

Mathematically, the patterns of harmonic modes of a dynamical system are estimated by the eigendecomposition of the Laplace operator. In the present case, the dynamical system of interest is the communication structure of the human brain, which is encoded in the dense functional connectivity (dense FC) matrix, in our study estimated from the pairwise temporal correlations between all pairs of vertices on the cortical surface (59.412 vertices in total; Figure 1a-c). We utilized the discrete counterpart of the harmonic modes defined on the graph built from the dense FC, i.e. the eigenvectors of the graph Laplacian (Figure 1d, e).

**Figure 1.**
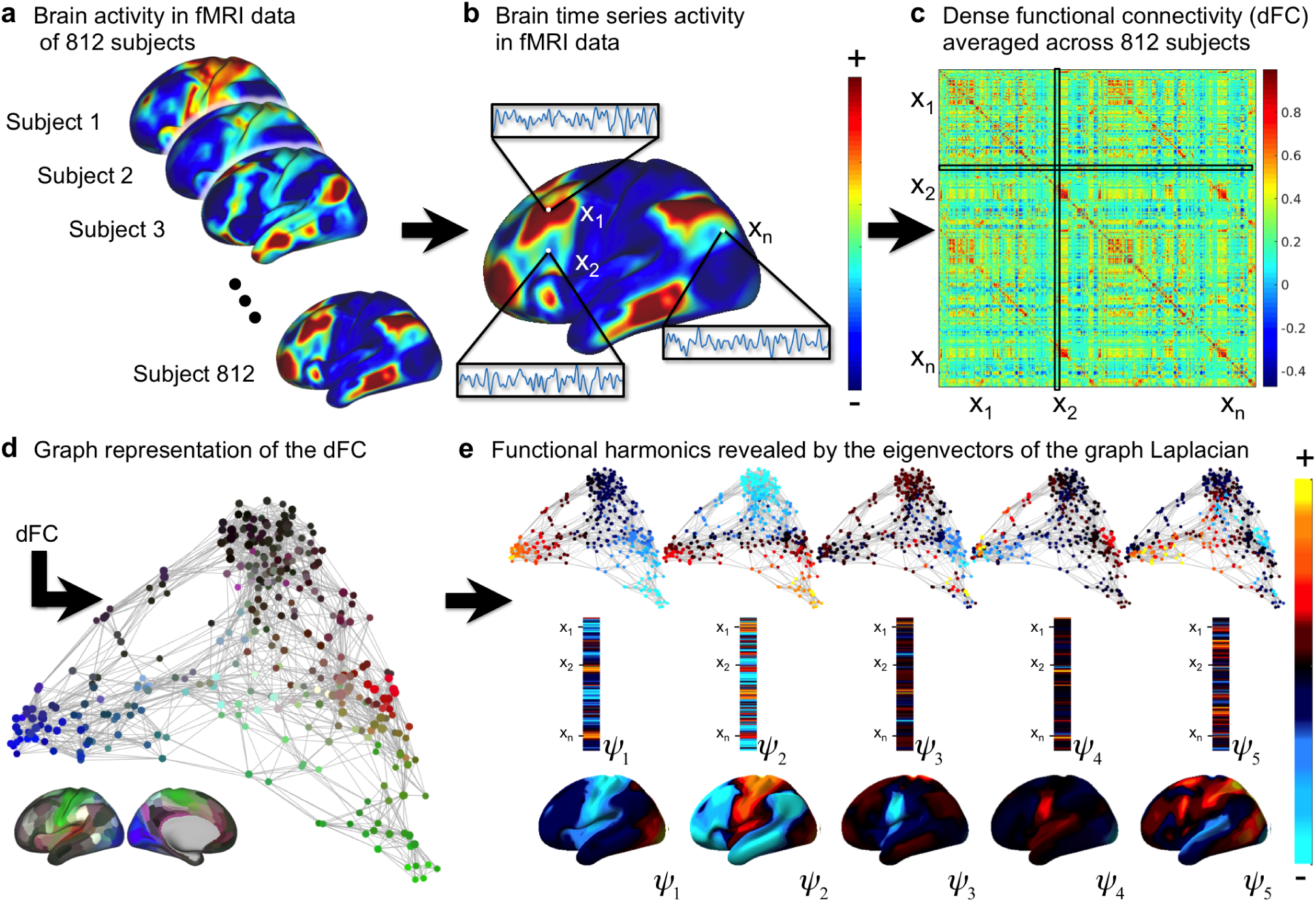
Workflow for the estimation of functional harmonics. **a**: Brain activity measured with functional magnetic resonance imaging (fMRI) in resting state for 812 subjects provided by the Human Connectome Project (HCP, 900 subjects data release).^41–48^ **b**: Illustration of brain activity time series of three representative vertices on the cortex (x_1_, x_2_, …, x_*n*_). **c**: The dense functional connectivity (FC) matrix computed from the temporal correlations between the time courses of each pair of vertices as shown in **b** averaged across 812 subjects. **d**: Representation of the dense FC as a graph, where the edges indicate strong correlations between the corresponding vertices. The anatomical location of the vertices are colour-coded^3^. **e**: Functional harmonics are estimated by the eigenvectors of the graph Laplacian computed on the graph representation of the FC. The first five functional harmonics ordered from the lowest to higher spatial frequencies are illustrated on the FC graph representation (top), in the eigenvector format as 59412 ×1 dimensional vectors (middle) and on the cortical surface (bottom). For illustrative purposes, the graph representations in **d** and **e** are shown for a parcellated version of the FC matrix using the HCP parcellation^3^.

Figure 2 shows the first 11 non-constant functional harmonics (referred to as [*ψ*_1_, *ψ*_2_, …, *ψ*_11_]), ordered starting from the lowest eigenvalue, and illustrating that each harmonic is a smoothly varying pattern on the cortex between a positive and a negative polarity; i.e., a gradient. There is an intrinsic relation between the Laplace eigenvalues and the spatial frequency/wavelength; namely as the eigenvalue increases, spatial frequency also increases, while the spatial wavelength decreases. Hence with increasing eigenvalue, the functional harmonics become increasingly more complex and segregate the cortex into an increasing number of nodal areas^40^ (contiguous areas of the cortex with similar colors in the surface plots in Figure 2). This intrinsic link between the harmonic frequency and cortical scale implies that functional harmonics yield not only a multi-dimensional, but also a *multiscale* description of the cortex. Note that the ordering by the wavelength/frequency is a property that emerges from the Laplace eigenfunctions and therefore is not present in function bases proposed by other methods, such as principle component analysis (PCA) or independent component analysis (ICA), which, as a result, do not implicitly possess this multiscale property.

**Figure 2.**
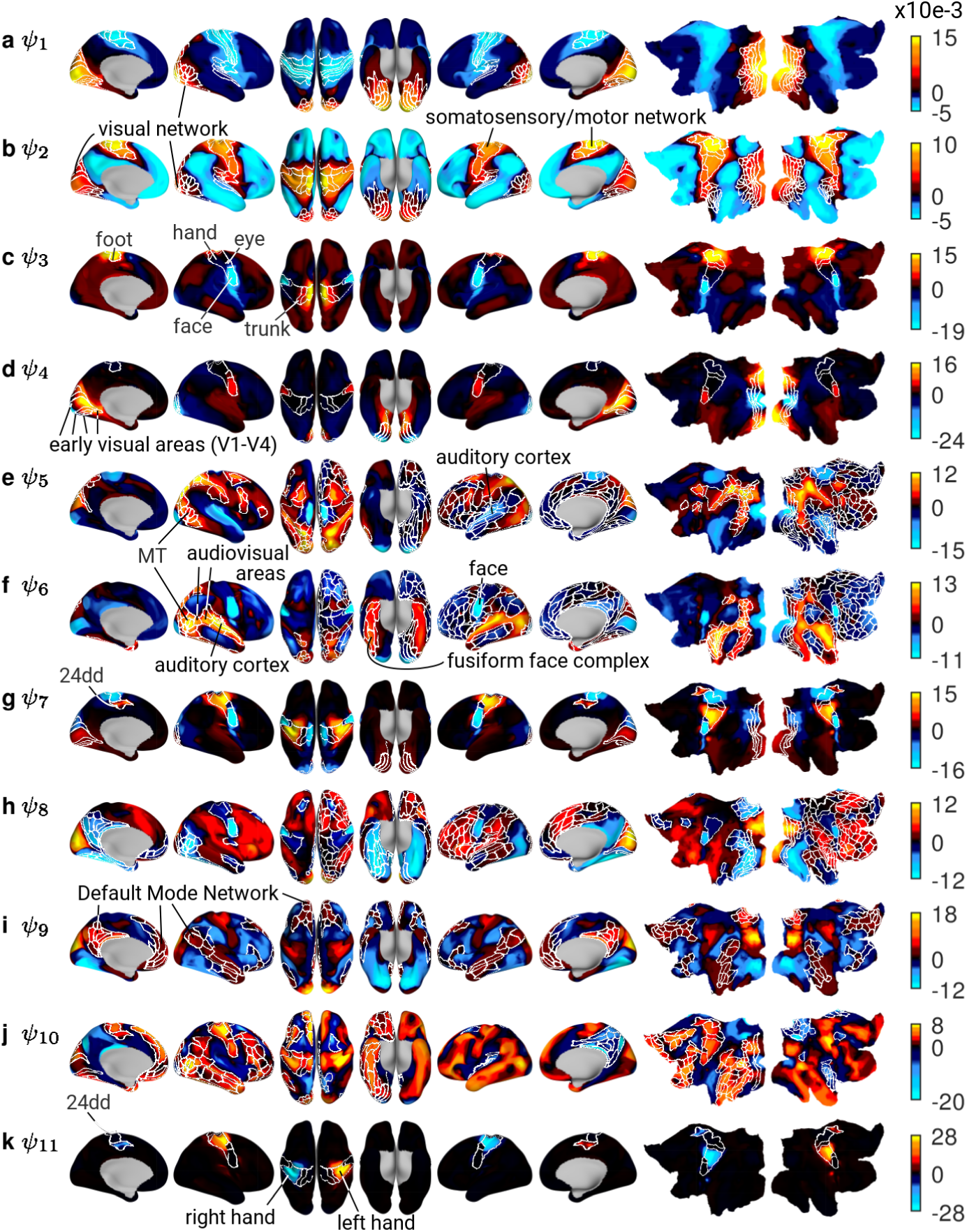
**a-k**: The first 11 non-constant functional harmonics plotted on the cortical surface. White lines show borders of HCP parcels. V1-V4: visual areas 1 to 4; MT: middle temporal visual area; 24dd: an area that contains a higher order representation of the hand; fusiform face complex: an area that responds specifically to images of human faces.

We note that the computation of the functional harmonics have been performed on the dense FC using 59412 *×* 59412 without using any parcellation.

### Functional harmonics capture sub-areal topographic organization

We first tested whether functional harmonics capture cortical organization on a sub-areal scale, i.e. within a cortical parcel. Specifically, we investigated whether functional harmonics capture somatotopy^15^ and retinotopy^14^, two major topographic mappings found in the brain. Topographic mappings represent sensory input on the cortical surface such that the relative positions of the receptors, which receive these inputs, are preserved, leading to a functional gradient *within* a specialized brain area.

Five somatotopic sub-areas (in each hemisphere), as defined by the HCP^3^, form a topographic map of the surface of the body on the cortex, i.e., the face, hands, eyes, feet, and trunk. We observed somatotopic mappings within functional harmonics 3 (*ψ*_3_), 7 (*ψ*_7_), and 11 (*ψ*_11_) (Figure 2c, g, k). Figure 3a illustrates the two-dimensional subspace formed by functional harmonics 3 (*ψ*_3_) and 11 (*ψ*_11_), which strikingly accounts for the precise mapping of the human body onto the somatotopic regions of the cortex (see SI Figure 1a-c for further examples). This example also illustrates how specialized functional regions emerge from the interaction of functional harmonics across multiple dimensions within the functional harmonics framework.

**Figure 3.**
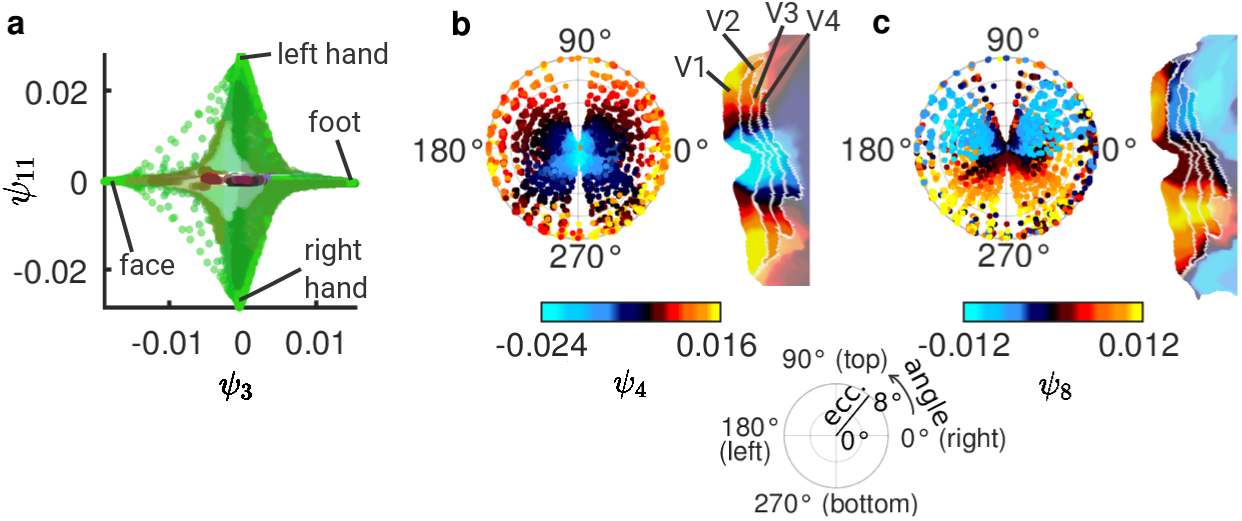
**a**: Functional harmonics 3 (*ψ*_3_) and 11 (*ψ*_11_) in their own space. Vertices are colour-coded according to their anatomical locations (see Figure 1d), and the location of 4 somatotopic areas in this space is annotated. **b** and **c**: Retinotopies of functional harmonics 4 (**c**; *ψ*_4_) and 8 (**d**; *ψ*_8_). Each panel shows, on the left, the colors of the respective functional harmonic in early visual areas V1-V4 on a polar plot of eccentricity (distance in degree from the fovea) and angle on the visual field (see legend at the bottom of the figure). On the right, the respective functional harmonic is shown on a flat map of early visual cortex (left hemisphere). V1, V2, V3, V4: visual areas 1, 2, 3, 4.

We then quantified the degree to which each somatotopic sub-area is delineated within these three functional harmonics by comparing the within- and between-area-variability specifically of the 10 somatotopic sub-areas (SI Figure 2a), thereby measuring their separation both from the rest of the cortex as well as from other somatotopic areas, indexed by a large silhouette value^49^ (see Methods for details). We compared the resulting values to those obtained from spherical rotations of functional harmonics 3 (*ψ*_3_), 7 (*ψ*_7_), and 11 (*ψ*_11_), in which we rotated the functional harmonic patterns on a spherical version of the cortical surface^50^. This control still yields smooth, symmetrical harmonic patterns on the cortex, but they do not emerge from the communication structure (FC matrix) of the brain and are not necessarily orthonormal. We found that for each of the tested functional harmonics, at least one somatotopic region is significantly separated (see SI Figure 2a; *p*_corr_ *<* 0.05 after Bonferroni correction, Monte Carlo tests with 300 rotated versions of the functional harmonics). This finding indicates that functional harmonics capture somatotopic organization in the cortex.

We next investigated the presence of retinotopic mapping of early visual regions (V1-V4), where cortical representations of the visual field reflect the positions of the receptors in the retina of the eye such that each vertex within the patterns of functional harmonics is assigned an eccentricity (distance from the fovea) and a polar angle (position in the visual field, i.e. top, bottom, left, right), according to the HCP retinotopy dataset^51^. Examples of polar plots of the retinotopic gradients are shown in Figure 3b, c (all polar plots are shown in SI Figure 2b). To investigate the degree of agreement between functional harmonics 1-11 (Figure 2) and the retinotopic mappings, we measured the correlation between eccentricity as well as polar angle maps and functional harmonic patterns 1-11 in V1-V4. We found significant correlations (*p*_corr_ *<* 0.05 after Bonferroni correction) between the retinotopic eccentricity map and all functional harmonics 1-11 except functional harmonic 9 (*ψ*_9_); and between the retinotopic angular map and functional harmonics 1-4 (*ψ*_1_, *ψ*_2_, *ψ*_3_, *ψ*_4_), 7-9 (*ψ*_7_, *ψ*_8_, *ψ*_9_), and 11 (*ψ*_11_).

These results demonstrate that retinotopic organisation of the early visual areas, which is normally extracted by applying retinotopic mapping techniques based on specific visual stimuli^14^, is implicitly present in the resting state brain activity and is revealed by the functional harmonic basis.

### Functional harmonics reveal specialized brain areas

As shown in Figure 2, the isolines of functional harmonics overlap with the borders of specialized brain regions (parcels) delineated by the HCP parcellation^3^ (all parcel borders are shown in SI Figure 3). In order to evaluate the degree of this correspondence, we quantified the overall overlap between functional harmonic patterns and brain regions (parcels). To this end, for each of the functional harmonics shown in Figure 2, we compared the within- and between-area-variability of each cortical region. A large difference between the within- and between-area variability of a particular region, indexed by a large silhouette value, indicates that that region is well-separated from the rest of the cortex^49^.

We compared our results to five alternative function bases. In order to test the effect of each step in our processing pipeline, we compared the performance of the functional harmonics first to that of the eigenvectors of the dense FC matrix (Figure 1c, SI Figure 4); second to the eigenvectors of the adjacency matrix (SI Figure 5), which is obtained after thresholding and binarizing the dense FC matrix, encoding the graph representation of human brain’s communication structure without involving the Laplace operator (Figure 1d); and third to a surrogate harmonic basis created by applying spherical rotations to the functional harmonics^50^. Furthermore, to relate the performance of functional harmonics to other well-known function bases, we also performed comparisons to the basis functions of PCA (SI Figure 6) and ICA (SI Figure 7).

**Figure 4.**
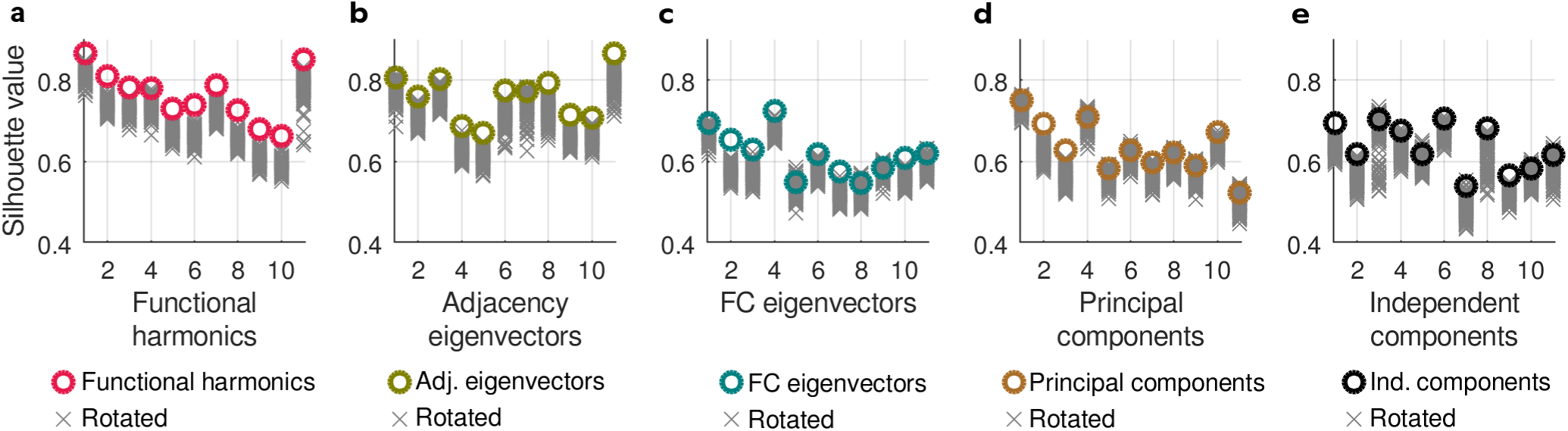
**a-e**: Silhouette values quantifying the degree to which isolines of the functional harmonics as well as control basis sets (coloured circles) and their rotations (gray crosses) follow the borders of the HCP parcellation. The silhouette value lies between -1 if all vertices were to be assigned to the wrong parcel, and 1 if all vertices were to be assigned to the correct parcel. For each basis function set, 220 rotations were computed.

**Figure 5.**
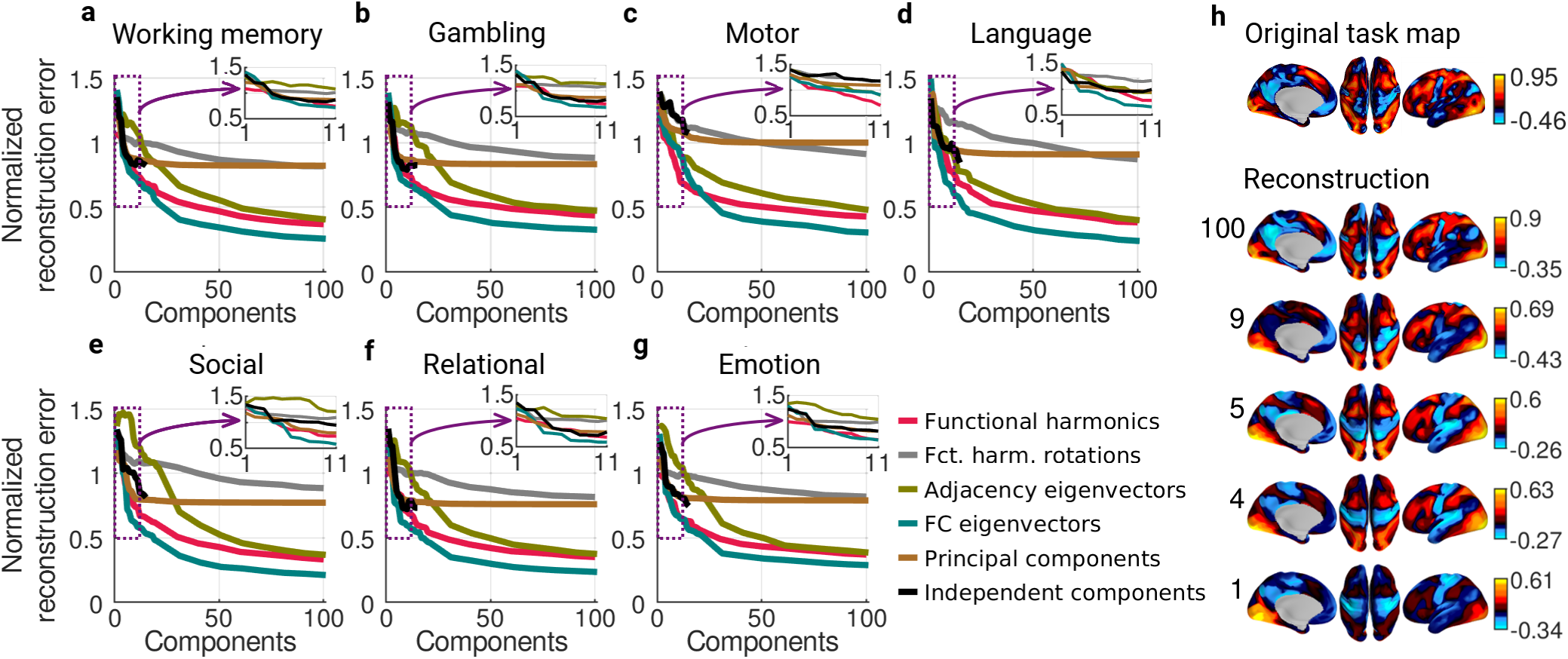
**a-g**: Mean reconstruction errors for each of the 7 task groups and all 6 basis function sets (see also SI Figure 9); **h**: One example for a reconstruction using a working memory task. The top panel is the original task activation map (working memory - body; see also SI Figure 10l), and subsequent panels use the number of harmonics indicated on the left to reconstruct it.

**Figure 6.**
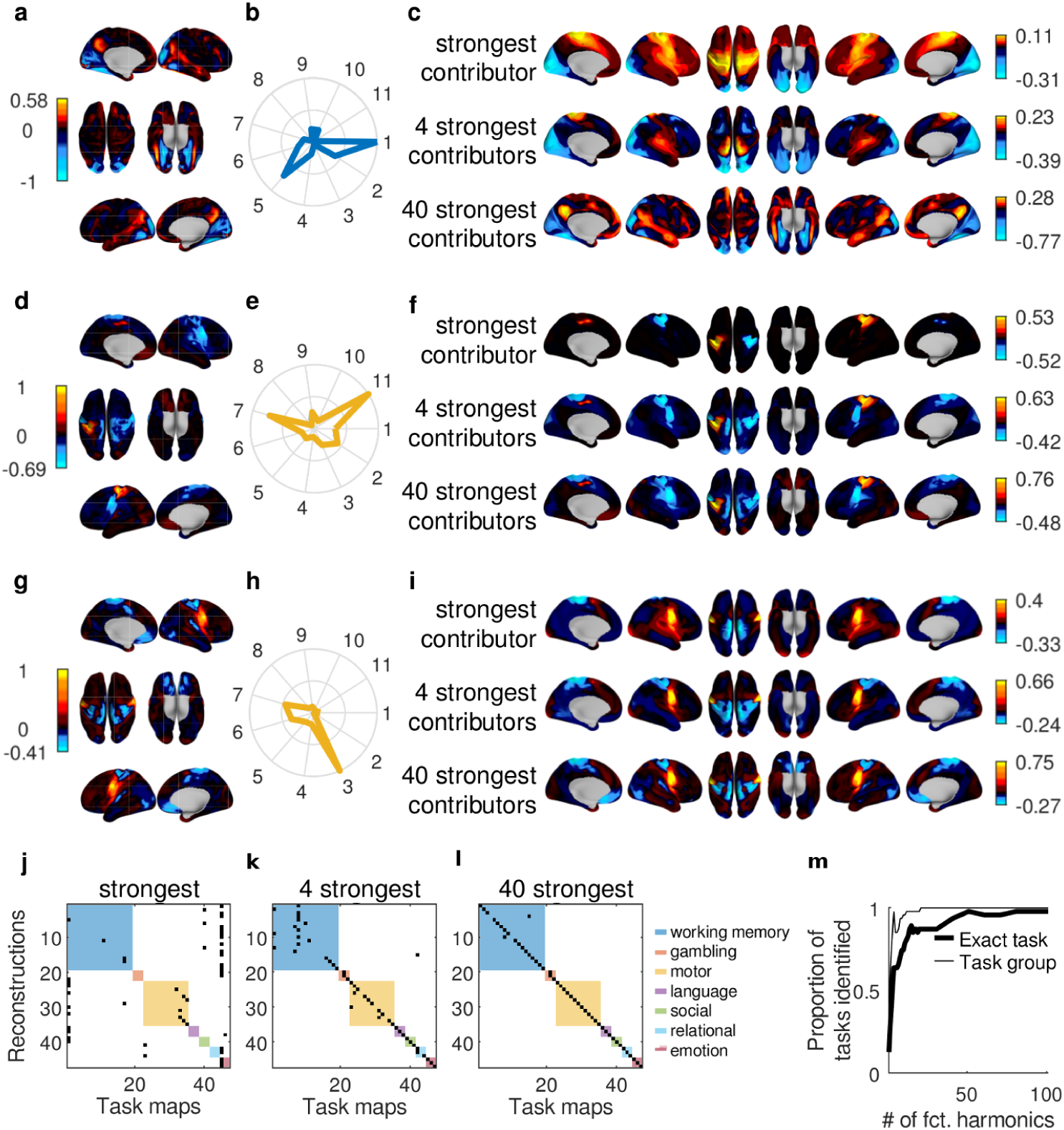
**a**: Map of the contrast between working memory (face) and average working memory from the HCP task dataset^56^, **b**: Contributions (normalized coefficients of the graph Fourier transform) for the first 11 non-constant functional harmonics, **c**: Reconstruction of the task map in **a** when using the functional harmonic with the strongest contribution (highest coefficient) only, the four functional harmonics with the strongest contributions, and the forty functional harmonics with the strongest contributions. **d-f**: The same as **a-c** using the map of the contrast between motor (right hand) and average motor. **g-i**: The same as **a-c** using the map of the contrast between motor (trunk) and average motor. **j-l**: Confusion matrices. Black entries mark the task map-reconstruction-pair which has the lowest reconstruction error, colored squares indicate the task group as in Figure 5. **m**: Proportion of reconstructions, for each number of harmonics, which have the minimum reconstruction error with their exact original task map (thick line) and a task map belonging to the same group of tasks as the original map (thin line).

As shown in Figure 4a, we found an alignment between the isolines of the functional harmonics and parcel borders for each of the first 11 functional harmonics, as verified by significantly larger silhouette values for functional harmonics compared to the rotated harmonic basis (*p*_corr_ *<* 0.05 after Bonferroni correction, Monte Carlo tests; see Materials and Methods for details). The only exception to this alignment was functional harmonic 4 (*ψ*_4_), which captures the retinotopic organization of early visual regions (see above for a discussion of retinotopic organization of functional harmonics). Importantly, this was not the case for any of the control function bases, where in each case at least some of the first 11 basis functions and their rotations performed equally well (Figure 4b-e). For qualitative evaluation, the overlap between parcels and functional harmonics as well as other bases are shown in SI Figures 3-7.

### Functional harmonics yield brain networks in frequency-specific communication channels

Another important aspect of the functional harmonics emerges due to their orthogonality: being an eigenvector of the Laplace operator, each functional harmonic is not only associated with a corresponding eigenvalue, related to its spatial frequency, but is also orthogonal to all other functional harmonics. This orthogonality implies that any functional harmonic is fully independent from all the other functional harmonics, thus providing a frequency-specific communication channel on the cortex.

In the following, we provide some insight into the functional significance of each of these communication channels, i.e. the functional harmonics shown in Figure 2. Functional harmonics 1 (*ψ*_1_) and 2 (*ψ*_2_) correspond to previously identified large-scale gradients^13^ that delineate the separation between the major sensory and the uni-vs. multimodal cortices in the brain, respectively (see SI Figure 1d). Figure 2a and b demonstrate the overlap between the visual and sensorimotor networks as defined in Yeo et al. (2011)^52^ and the gradiental patterns of the first and second functional harmonics. We observed that functional harmonic 3 (*ψ*_3_) reveals a finer subdivision of the somatosensory/motor system^53–55^. The overlay of the borders of the five somatotopic areas defined by the HCP^3, 56^ on the third functional harmonic are shown in Figure 2c. Similarly, in functional harmonic 4 (*ψ*_4_), we found a finer segregation of the visual system, following a retinotopic eccentricity gradient^51^. The overlay of the borders of early visual areas (V1-V4) on functional harmonic 4 (*ψ*_4_) are shown in Figure 2d (for further details on retinotopic and somatotopic mapping, see above). These results clearly demonstrate that each of these functional networks previously identified in the literature, simply correspond to a different communication channel of the human brain, occurring in parallel in the multi-dimensional functional harmonic representation.

Qualitative evaluation of higher frequency functional harmonics systematically revealed their link to more specialized complex brain function. The pattern observed in functional harmonic 5 (*ψ*_5_, Figure 2e) is consistent with a communication channel in which action and perception interact. In the negative polarity, we found primary visual, auditory, and somatosensory cortices, while the regions in the positive polarity closely resemble the sensory-motor pathway, which has been shown to mediate selective interactions between resting state networks along the visual hierarchy^52^. Parts of this pathway are known to be modulated by visuospatial attention^57^. The fact that we also found auditory and somatosensory regions is in line with the idea that the interplay between action and perception circuits - also known as active inference^58, 59^ - is a multimodal process^60–62^. In functional harmonic 6 (*ψ*_6_), auditory and visual areas were both localized in the positive polarity, forming a network related to audiovisual object (including faces) recognition^63–65^, i.e. recognition of the “outer world”. The negative polarity of functional harmonic 6 (*ψ*_6_) segregates the somatotopic face area as well as parts of the default mode network (DMN), a network of regions whose activity has been related to self-referential tasks^66^. Thus, the negative polarity of functional harmonic 6 (*ψ*_6_) forms a self-referential processing stream^66–68^. Functional harmonic 7 (*ψ*_7_) provides a finer somatotopic gradient, including a higher hand area, 24dd, in the medial cortex^53^ (see Figure 2g and annotations in Figure 2c). Functional harmonics 8 to 10 (*ψ*_8_, *ψ*_9_, *ψ*_10_) correspond to different subdivisions of higher order networks such as the frontoparietal network and DMN (see SI Figure 8a-c). In particular, the DMN^69^ is delineated in the positive polarity of functional harmonic 9 (*ψ*_9_) (borders of the DMN as defined by Yeo et al. (2011)^52^ are overlaid on functional harmonic 9 (*ψ*_9_) in Figure 2i). In functional harmonic 10, we found a significant correlation (*r* = *–*0.63, *p* = 4 *·* 10^*–*21^) with the degree of auditory involvement of the functional areas (SI Figure 8d). Functional harmonic 11 (*ψ*_11_), the first clearly asymmetric harmonic between the two hemispheres, yields the separation between the right and left somatotopic hand areas^70^.

Overall, these results demonstrate that functional harmonics provide a multitude of functionally relevant communication channels, each associated with a unique spatial frequency, and enable a set of parallel processing streams in the human brain.

### Functional harmonics provide the basis functions of human cognition

Given our results demonstrating how highly complex function of the human brain can simply be orchestrated by simultaneous processing across different functional harmonics, we hypothesized that they can provide the building blocks of complex human behaviour. Remarkably, the implicit link between the functional harmonics and the well-known Fourier basis provides the very needed property for them to constitute a language to describe any pattern of brain activity. The Laplace eigenfunctions on a one-dimensional domain with cyclic boundary conditions, i.e. a circle, yield the sine and cosine functions with different frequencies, which comprise the Fourier basis. Thus, functional harmonics being the eigenfunctions of the Laplace operator applied to the dense FC, provide the extension of the Fourier basis to the communication structure of the human brain. As such, they provide per definition a frequency-specific function basis, in which any pattern of brain activity can be represented as a weighted combination of functional harmonics. Given the experimental evidence showing that resting state functional connectivity reflects connectivity during task^9–12^, we tested how efficiently the functional harmonics can represent the task-induced brain activity measured on the cortex. To this end, we reconstructed 47 group-level task maps provided by the HCP^56^ from the superposition of functional harmonics (see Materials and Methods). The 47 maps consist of activation maps as well as contrasts derived from 7 groups of tasks (working memory, motor, gambling, language, social, emotional, relational - see Materials and Methods for summaries). The functional harmonic reconstruction yields a coefficient (weight) for each functional harmonic, quantifying how much it contributes to a certain task map. The set of all coefficients forms a spectrum equivalent to the power spectrum obtained from a Fourier transform, in this case the power spectrum of the functional harmonic basis.

We first tested whether it is possible to approximate task maps as superpositions of subsets of functional harmonics, linearly combining them in the order of their eigenvalues. We quantified the goodness of fit by measuring the distance between the original and the reconstructed task maps. Figure 5a-g show the average normalized reconstruction errors (i.e., the original and reconstructed task maps were normalized to zero mean and unit variance in order to ensure comparability across tasks) for all groups of tasks and for all compared function bases: for the functional harmonic basis (red line), the error drops from between 1.00 (for Emotion) and 1.40 (for Language) to between 0.65 (for Emotion) and 0.78 (for Language) when using only the first 11 functional harmonics shown in Figure 2, which constitute 0.02% of the total functional harmonic spectrum. This corresponds to a level of correlation between the original task maps and the task maps reconstructed from the first 11 non-constant functional harmonics of between 0.78 (for Emotion) and 0.69 (for Language; see also SI Figure 9a). Figure 5h illustrates the reconstruction procedure for one specific task (working memory: body; see SI Figure 10 for all tasks).

We compared the performance of the first 11 non-constant functional harmonics in reconstructing task activation maps to that of the control function bases (rotations of functional harmonics, eigenvectors of the FC, eigenvectors of the adjacency matrix, principal components, and independent components; Figure 5a-g, SI Figure 9b-e). We found that functional harmonics outperform the rotated harmonic basis (*p*_corr_ *<* 0.005, Monte-Carlo tests with 1000 permutations, Bonferroni corrected for multiple comparisons), adjacency eigenvectors (*p*_corr_ *<* 0.005, Monte-Carlo tests with 1000 permutations, Bonferroni corrected for multiple comparisons), as well as PCA (*p*_corr_ *<* 0.005, Monte-Carlo tests with 1000 permutations, Bonferroni corrected for multiple comparisons) and ICA (*p*_corr_ *<* 0.005, Monte-Carlo tests with 1000 permutations, Bonferroni corrected for multiple comparisons), and did not exhibit any significant difference to the performance of the FC eigenvectors (*p >* 0.15 before correction for multiple comparisons, not significant (n.s.)).

In order to examine the reconstruction performance of each function basis for different task groups, we applied the same Monte-Carlo analysis to each of the 7 task categories separately. We found that reconstruction errors of functional harmonics were significantly lower than those of their rotations for each of the task groups (all *p*_corr_ *<* 0.035, Monte-Carlo tests with 1000 permutations, Bonferroni corrected for multiple comparisons), and significantly lower than those of the adjacency eigenvectors in six out of seven task groups (all *p*_corr_ *<* 0.035, Monte-Carlo tests with 1000 permutation, Bonferroni corrected for multiple comparisons, except language, where *p* = 0.18, before correction for multiple comparisons, n.s.). In comparison to FC eigenvectors, while there was no significant difference in the reconstruction performance when all tasks were pooled, we found that functional harmonics performed significantly better in the reconstruction of motor tasks (*p*_corr_ *<* 0.035, Monte-Carlo tests with 1000 permutations, Bonferroni corrected for multiple comparisons; see inset in Figure 5c). Compared to PCA and ICA, the reconstruction errors of functional harmonics were significantly lower for motor and working memory task groups (all *p*_corr_ *<* 0.035, Monte-Carlo tests with 1000 permutation, Bonferroni corrected for multiple comparisons), while for all other task groups there were no significant differences (all *p >* 0.01 before correction for multiple comparisons, n.s.). These results indicate that functional harmonics delineate the functional systems involved in working memory and motor tasks more precisely than other function bases used as control. It is important to note that the number of tasks in the remaining categories is smaller (3 tasks per category) than that of the motor and working memory task groups, and more data may be required to achieve significant differences for these categories. In summary, when all individual task groups as well as the overall performance in reconstructing the complete task pool is considered, the functional harmonics outperform all 5 control function bases using only the first 11 non-constant components.

Given that functional harmonics constitute functionally relevant communication channels, we hypothesized that the task activation maps can be characterized by their power spectrum. Figure 6a, d, g and Figure 6b, e, h show two examples of task activation maps and the corresponding normalized power of the first 11 non-constant functional harmonics, respectively, revealing how strongly each of the 11 functional harmonics shown in Figure 2 contributes to these particular task maps. For qualitative evaluation, we display the task activation maps reconstructed by superimposing functional harmonics in the order of their contribution strength for varying numbers of functional harmonics in Figure 6c, f, i (see SI Figure 10 for all tasks). Across all 47 task maps that were evaluated, the functional harmonic which was the strongest contributor was always either the constant functional harmonic or one of the first 11 non-constant harmonics shown in Figure 2.

In order to evaluate the uniqueness of the functional harmonic power spectrum of each task activation map, we computed the distance between a given reconstructed map and all original task maps, resulting in a confusion matrix for each number of harmonics with maximum contribution. If task maps can indeed be characterized by their functional harmonics power spectra, the error should be minimal between a reconstruction and its corresponding task map compared to the error of the reconstruction of the other 46 task maps. The confusion matrices in Figure 6j-l show the pairs of the original and reconstructed task activation maps with the minimum distance when using 1, 4, and 40 functional harmonics with maximum contribution. Coloured squares mark the 7 task groups as in Figure 5. The proportion of unambiguously identified tasks in relation to the number of functional harmonics is shown in Figure 6m. We found that sparse representations using the 4 functional harmonics with the largest power for each task are sufficient to unambiguously characterize the seven task groups with the exception of one working memory task (Figure 6k), and 70% of all individual tasks. When the 40 functional harmonics with maximum contribution are used, which corresponds to 0.1% of the complete spectrum of functional harmonics, 44 out of 47 task maps are correctly identified from their reconstructions (Figure 6l).

These results demonstrate that that functional harmonics provide a novel functionally relevant representation, where the brain activity accompanying different tasks can be uniquely identified from the activation profiles of a small range of functional harmonics.

Overall, our findings show that functional harmonics not only correspond to a multitude of the functionally relevant networks of the human brain observed in resting state, but also effectively reconstruct task-induced brain activity, thus suggesting that they provide a strong candidate for the fundamental building blocks of higher order complex human brain function and behaviour.

## Discussion

We reveal a previously unknown principle of cortical organization by applying a fundamental principle ubiquitous in nature - harmonic modes - to the communication structure of the human brain. The resulting modes termed *functional harmonics* link observations spanning several spatial scales, from sub-areal topographic mappings (somatotopy, retinotopy), emergence of specialized functional brain areas, to the level of large-scale networks and combinations thereof. In this view, functional integration and segregation on all these levels are organized according to the same underlying principle, i.e., harmonicity.

In this framework, functional networks observed during rest and task emerge as a data-driven, frequency-specific function basis derived from the human resting state functional connectivity matrix. We demonstrate the meaning of the first 11 functional harmonics as functional networks of the human brain. Functional harmonics are estimated as the eigenvectors of the graph Laplacian. As such, they are separated by spatial frequency and orthogonal to one another, making it possible to interpret them as communication channels that provide an orthogonal function basis. Any pattern of cortical activity can be expressed in this function basis as a superposition of functional harmonics. In this new function basis, brain activation patterns measured during tasks are understood as task-specific spectra quantifying the contribution of each functional harmonic, allowing different combinations of the same functional networks to be active in order to fulfill task demands. Importantly, functional harmonics and the networks to which they correspond, emerge in a wholly self-organizing fashion from the functional connectivity.

The brain has to be able to rapidly switch between rest and different task demands. Functional harmonics are, by definition^71^, the smoothest patterns that respect the constraints posed by the functional relationships given by the FC. This implies that the average difference between neighbouring nodes in a graph representation is minimized. Intriguingly, theoretical work has shown that activation patterns on graphs in which neighbouring nodes co-activate lead to patterns with minimum free energy or entropy^72–74^, and that the transition between such patterns requires minimal energy^75^. This means that transitioning between the functional networks instantiated by the functional harmonics in response to changing task demands is optimally efficient.

Here, we explicitly show that functional harmonics are building blocks of cognitive activity in the brain by characterizing a multitude of task activation maps from their functional harmonic reconstructions. In particular, our results demonstrate that although there is a multitude of function bases one can choose to represent patterns of brain activity such as the well-known principal components or independent components of PCA and ICA, functional harmonics stand out in their ability to capture certain aspects of cortical organization: our findings reveal that out of the 5 function bases used to represent patterns of cortical activity; i.e. *(i)* eigenvectors of the FC matrix, *(ii)* eigenvectors of the adjacency matrix, *(iii)* rotated versions of functional harmonics, *(iv)* PCA and *(v)* ICA, only the functional harmonics yield both, a delineation of cortical areas *and* an efficient reconstruction of task activation maps, and thus provide the strongest candidate to be the basis functions of human cognition. Harmonic function bases are set apart from the other bases that we tested by the fact that basis functions exhibit an implicit ordering according to their wavelength (spatial frequency), and hence provide not only a multi-dimensional but also a multiscale representation of brain activity. Our results suggest that this ordering captures an important aspect of cortical organization, as functional harmonics are able to outperform each of the alternative function bases in terms of how well the parcellation is captured and how well task activation patterns are reconstructed.

Moreover, functional harmonics unify the competing views that brain activity arises *either* from smoothly varying gradients *or* from the modular and specialized regions. Within the functional harmonic framework, specialized regions correspond to isolines of the gradiental patterns of functional harmonics, where each region can also be increasingly segregated into finer regions through the interaction of functional harmonics across multiple dimensions. Hence our findings provide, to our knowledge, the first principle that unifies the gradiental and modular aspects and reveals the multi-dimensional nature of cortical organization.

Considering that the principle of harmonic modes when applied to the structural connectivity of the human brain - the human connectome - have been shown to reveal the functional networks^36^, our results point to the emergence of the same fundamental principle in multiple aspects of human brain function. Beyond the results presented here, functional harmonics suggest novel ways to understand the dynamics of the human brain in health and in pathology as well as to explore individual differences within this multi-dimensional harmonic representation.

## Materials and Methods

### Data

The data used in this study was acquired and made publicaly available by the Human Connectome Project, WU-Minn Consortium (Principal Investigators: David Van Essen and Kamil Ugurbil; 1U54MH091657) funded by the 16 NIH Institutes and Centers that support the NIH Blueprint for Neuroscience Research; and by the McDonnell Center for Systems Neuroscience at Washington University. All study protocols were approved by the Washington University institutional review board, and informed consent was obtained in all cases^41, 42^.

In this study, we used the dense functional connectivity (FC) matrix, which is part of the Human Connectome Project’s 900 subjects data release^41–48^. It is available under db.humanconnectome.org/data/projects/HCP 1200^76^. Clicking on “812 Subjects, recon r227, Dense Connectome” will download the appropriate .zip-archive (user login necessary). The list of names of all the files used in this study is shown in Table 1. Note that in this release, many of the subjects are related to at least one other subject of the group. The group average functional connectivity matrix was obtained by correlating group-PCA eigenmaps from 812 out of the 900 subjects included in this release, which are the subjects that having completed all four sessions of 15-minute resting state fMRI.

**Table 1.**
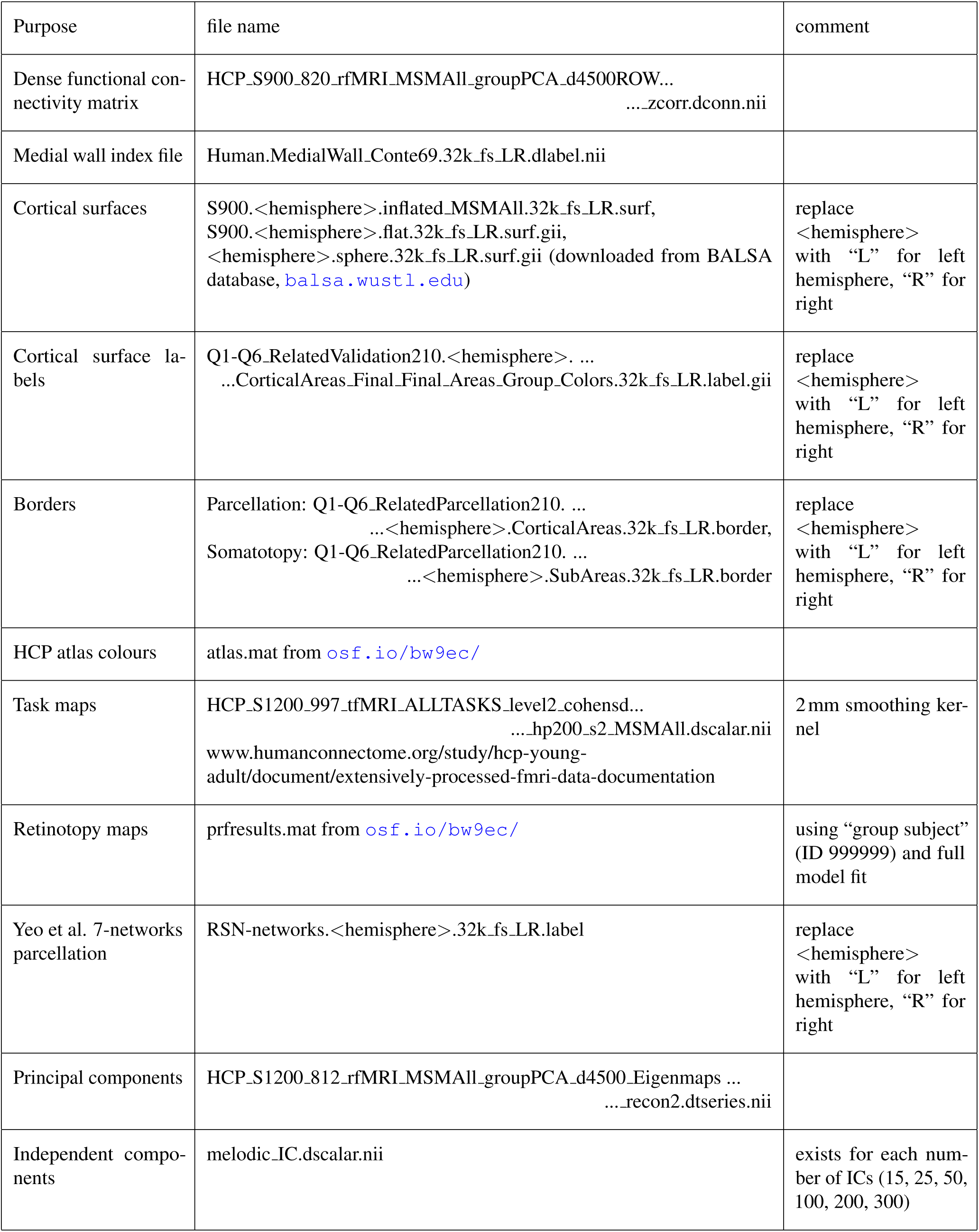
Files used in our computations. All data was downloaded from the human connectome project database (db.humanconnectome.org/data/projects/HCP 1200) unless otherwise specified.

For task reconstructions, we used data contained in the S1200 group average data release, which is available on www.humanconnectome.org/study/hcp-young-adult/document/extensively-processed-fmri-data-documentation, as “HCP S1200 GroupAvg v1 Dataset”.

For the analyses involving retinotopic maps, we used data available on osf.io/bw9ec/ and described in Benson et al. (2018)^51^. The relevant file is named “prfresults.mat” and contains a variable “allresults” of dimensionality 91282 (grayordinates) × 6 (quantities) × 184 (181 subjects plus 3 different group averages) × 3 (model fits). We used only the quantities ‘ang’ and ‘ecc’, the first model fit, of the group average across all available subjects, which uses all available time points. See osf.io/bw9ec/wiki/home/ for details.

Data are encoded in CIFTI file format^41^, which means that coordinates are defined on the cortical surface (“grayordinates”), i.e. using *n* vertices rather than voxels^48^. The file was read using connectome workbench functions^76^ and converted to a single precision vector of length (*n · n* – *n*)*/*2 (due to its symmetry) using Matlab^77^. We also excluded the medial wall. This reduced the size of the FC matrix in memory from 33 GB to approximately 6 GB, greatly easing subsequent computations. The loss in precision is negligible compared to the accuracy with which pairwise correlation can be estimated from noisy fMRI time courses.

For visualization purposes, we used the surfaces provided with the functional data.

### Software

All data analysis was performed using MATLAB 2014b or 2017b, using also scripts and functions from the following freely available software packages:

- Fieldtrip version 20180903
- Connectome workbench (https://www.humanconnectome.org/software/connectome-workbench)
- gifti toolbox (https://www.artefact.tk/software/matlab/gifti/)

### Background: Functional Harmonics

The approach presented here relies on representing the human brain’s communication structure (dFC) as a graph and estimating the eigenfunctions of graph Laplacian applied to this structure. The graph representation of the brain’s communication structure *𝒢* = (*𝒱, ℰ*) is created by representing the vertices sampled form the gray matter cortical surface as the nodes *𝒱* = {*v*_*i*_ |*i* ∈1,, *n*} with *n* being the total number of nodes (*n* = 59.412 in this study) and by representing the connections between the vertices as the edges *ℰ* = {*e*_*i j*_ |(*v*_*i*_, *v* _*j*_) ∈ *𝒱* × *𝒱*}, which come from the connections in the dFC matrix. We represent this graph structure *𝒢* by its *n×n* adjacency matrix *A* = [*a*_*i j*_] that is formed by connecting each node *i* to its *k*-nearest neighbours (*k* = 300 in this study) according to its correlations in the dFC matrix, i.e.:

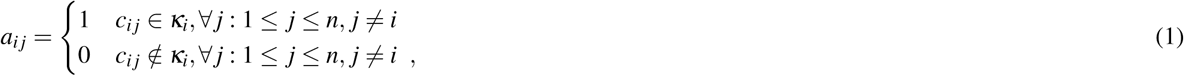

where *κ*_*i*_ is the set of the *k* largest values in row *i* in the dFC matrix. In order to ensure *A* is symmetric, we also set *a* _*ji*_ = 1, if *a*_*i j*_ = 1. Defining **A** as such results in a symmetrical sparse binary matrix.

Then we estimate the graph Laplacian defined as

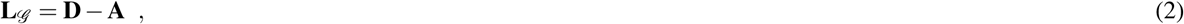

where **A** is the adjacency matrix as defined above, and **D** is the degree matrix, which is defined as a diagonal matrix with diagonal elements

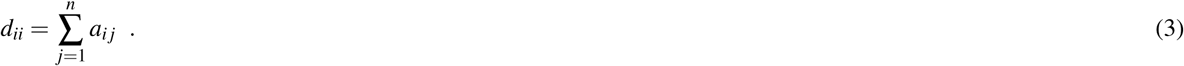

As such, the degree matrix **D** contains each node’s degree in its diagonal. Finally, we estimate the functional harmonics as the eigenfunctions Ψ = *{ψ*_1_, *ψ*_2_, *…, ψ*_*n*_*}* by solving:

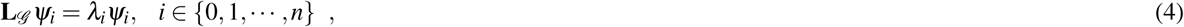

where *ψ*_*i*_ are the *n ×* 1 eigenvectors and *λ*_*i*_ are the corresponding eigenvalues.

### Control function bases

1. Spherical rotations: We performed comparisons against spherical rotations of surface maps. We followed^50^, adapting freely available code (github.com/spin-test/spin-test) to be used with HCP surfaces. In this approach, surface maps are projected to a spherical surface and then rotated by a random angle. Values are then mapped back to the nearest vertex, and the map is symmetrized in order to preserve this property. Parts of the corpus callosum that are rotated to the cortical surface are labelled as missing data (NaNs) and are ignored in any subsequent calculations (e.g. within- and across area distances, see below). Since we used multi-dimensional function based, we rotated the surface maps corresponding to each dimension by the same angle. Note that, however, the resulting rotated function basis is no longer orthonormal due to the symmetry preserving step.
2. Eigenvectors of the dense FC: An intuitive basis is to take the eigenvectors of the dense FC without applying a threshold as done for obtaining the adjacency matrix. These eigenvectors have been shown to contain valuable information about dynamical FC^78^. The first 20 eigenvectors of the dense FC are shown in SI Figure 4.
3. Eigenvectors of the adjacency matrix: In order to test the effect of thresholding/binarizing on the one hand and the effect of using the graph Laplacian instead of the adjacency matrix itself on the other, we also compared to the eigenvectors of the adjacency matrix, i.e. the dense FC thresholded such that only the 300 nearest neighbors of each vertex are retained and set to 1. The first 20 eigenvectors of the adjacency are shown in SI Figure 5.
4. Principal components (PCs): PCA (principal component analysis) is a popular dimensionality reduction technique which preserves the maximum amount of variance in the data. It consists of taking the eigenvectors of the covariance matrix of the time series. These principal components are provided by the HCP via Connectome DB (see Table 1). The first 20 PCs are shown in SI Figure 6.
5. Independent components (ICs): A very popular dimensionality reduction technique in resting state fMRI^79^, independent component analysis is the foremost method for obtaining resting state networks. It consists of analyzing the time series of the data and finding those spatial patterns that are maximally independent. We tested all sets of ICs that are provided by the HCP (see Table 1, and found that the set with the lowest number of components, i.e. *n* = 15, performs best. Therefore, we restricted our comparisons to this set of ICs. Note that ICs are not orthonormal and therefore do not form a basis in the strictly mathematical sense. The 15 ICs used in our comparisons are shown in SI Figure 7.

### Monte Carlo simulations

We used a Monte-Carlo approach for statistical validation.

For the silhouette values, we followed^50^, where permutations consist of rotated surface maps (see previous section) of the functional harmonics as well as principal components, independent components, eigenvectors of the dense FC, and eigenvectors of the adjacency matrix. Silhouette values were then computed for the original, non-rotated map as well as for *n* = 220 rotated maps, and p-values were computed based on the number *P* of rotations that performed better than the original map:

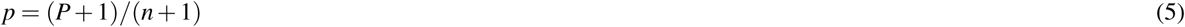

We performed Bonferroni correction by multiplying the resulting p-value by 11, i.e. the number of dimensions that was tested.

We used the same approach for the somatotopy index, but only applied to the functional harmonics and their rotations. Since in this case, we had five somatotopic areas (we averaged over the two hemispheres) and tested three of the 11 functional harmonics (*ψ*_3_, *ψ*_7_, and *ψ*_11_), we required *n* = 300 rotations in order to achieve a significance level of *α* = 0.05 with 15 comparisons.

We also applied a Monte-Carlo permutation test to the mean reconstruction errors by permuting the labels of the basis 1000 times for each control basis. Here, we pooled the reconstruction errors over the first 11 non-constant components. For the overall reconstruction performance, we also pooled all 47 task maps; for ad-hoc tests of each task category, we pooled only over the tasks in each category.

### Silhouette values

To test whether isolines of the functional harmonics follow the boundaries of the parcels as defined in the HCP parcellation^3^, we compute the silhouette value^49^ of each functional harmonic as:

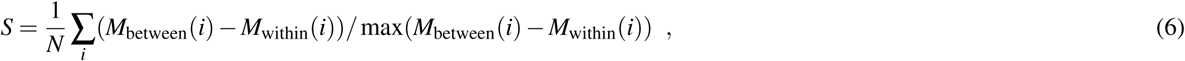

where *M*_between_(*i*) is the average Euclidean distance between vertices belonging to a parcel *i* and vertices belonging to all other parcels, while *M*_within_(*i*) is the average distance between vertices within the parcel *i*. If all vertices belonging to a parcel *i* have the same value, and at least some vertices outside the parcel *i* have different values, then *M*_between_(*i*) *>* 0, *M*_within_(*i*) = 0 and *S*(*i*) = 1. By averaging over the silhouette values of all parcels, one obtains a measure of how well the data fit the parcellation. Note that we replaced the somatosensory/motor core areas 1, 2, 3a, 3b, and 4 with the somatotopic sub-areas given by the HCP^3^ for a more detailed evaluation.

To evaluate the somatotopic organization of the functional harmonics, we use a measure that was similar to the silhouette value, but adapted to measure the separation from the rest of the cortex *and* from other somatotopic areas.

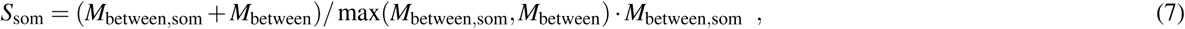

where *M*_between,som_ is the average Euclidean distance between vertices belonging to a somatotopic area and all other vertices belonging to all other somatotopic areas. The first term of the equation is between 1 and 2 and is close to 2 if both *M*_between,som_ and *M*_between_ are equal. Multiplying by *M*_between,som_ ensures that *S*_som_ is not large if both *M*_between,som_ and *M*_between_ are small.

### Task maps

We used group-averaged task activation maps provided with the S1200 group average data release of the HCP (see Table 1, www.humanconnectome.org/study/hcp-young-adult/document/extensively-processed-fmri-data-documentation). Here we provide a summary of the tasks that form part of the HCP task battery^56^. There are 7 groups of tasks: working memory, motor, gambling, language, social, emotional, relational. Subjects performed all tasks in two separate sessions (working memory, gambling, and motor in the first session, language, social cognition, relational processing, and emotion processing in the second).

#### Working memory

Four different stimulus types were used, presented in separate blocks: pictures of faces, places, tools and body parts. Two different task types were used: a 2-back working memory task, where subjects had to respond if a stimulus matched that two trials back, and a 0-back working memory task, where subjects had to respond whenever a single stimulus returned that was presented at the beginning of the block. This results in a total of 19 different working memory task maps, consisting of 14 activation maps (such as 0-back, 2-back, face, body, etc.) and 5 contrasts (between the two task types, between each stimulus type and the average across all stimuli, etc.).

#### Motor

Visual cues indicated whether participants should move their left or right fingers, left or right toes, or move their tongue. The goal was to identify the motor areas that correspond to these five body parts. This results in 26 different task maps (7 activation maps for 5 body parts plus visual cue plus average, and 6 contrast maps).

#### Gambling

(Incentive processing.) Subjects played a game in which they could win or lose money. The game was to guess whether the number on a “mystery card” that could range between 1 and 9 would be less or more than 5. The numbers were given after subjects made their guess and were chosen according to the trial type: “win” - the number would correspond to their guess and they would win 1$; “neutral” - the number would equal 5 and they would neither win nor lose any money; “loss” - the number would not correspond to the guess and participants would lose $0.50. Separate blocks are used in which trials are either mostly win or mostly lose, resulting in two conditions, punish and reward. This results in 3 different task maps (2 activation maps, i.e. one for each condition, and 1 contrast).

#### Language

Two different task types were used, “story” and “math”. “Story” consisted of participants listening to 5-9 sentences of a story, and answering a 2-alternative forced choice question thereafter. “Math” required participants to solve simple addition and subtraction problems. The two task types are similar in terms of auditory input and attentional load, but different in terms of semantic and numerosity related processing. As for gambling, the two task types result in 3 task maps (2 activation, 1 contrast).

#### Social

(Theory of Mind, TOM.) Subjects viewed videos of objects (squares, circles, triangles) that moved around in one of two ways: “Random” - there was no interaction between the objects, or “TOM” - the objects moved as if they were reacting to the other objects’ “thoughts and feelings”. They then had to judge whether the objects were interacting or not, or respond with “not sure”. As with gambling and language, the two task types result in 3 task maps (2 activation, 1 contrast).

#### Emotional

Subjects viewed one of two types of stimuli, “faces” or “shapes”, and had to decide which of two stimuli presented at the bottom of the screen matched the stimulus at the top of the screen. The faces included emotional stimuli, i.e. angry or fearful expressions. Again, the two task types result in 3 task maps (2 activation, 1 contrast).

#### Relational

There were two conditions, “match” and “relational”. In all cases, stimuli can have one of six shapes combined with one of six textures. In the “match” condition, which served as a control condition, two shapes were presented at the top and one at the bottom of the screen. A word (“shape” or “texture”) that appears in the middle of the screen instructs subjects to decide whether the bottom stimulus matches either of the top stimuli in the dimension indicated by the word. In the “relational” condition, two stimuli are presented each at the top and at the bottom of the screen, with no word in the middle. Instead, participants have to determine themselves across which dimension the top pair differs, and, subsequently, indicate whether the bottom pair differs over the same dimension. Again, the two task types result in 3 task maps (2 activation, 1 contrast).

Task maps were computed using FSL’s FEAT and FLAME^80, 81^ and conducting a between-subject (“level 2”) analysis, resulting in effect sizes (Cohen’s d). We used the task maps with minimal smoothing (2mm total smoothing); see 1200 subjects data release reference manual, pp. 45-54 and 100-104.

### Reconstructing the task maps from functional harmonics

The spatial pattern of each task map on on the cortex **s**(*v*) was decomposed into and reconstructed from the functional harmonics 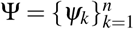

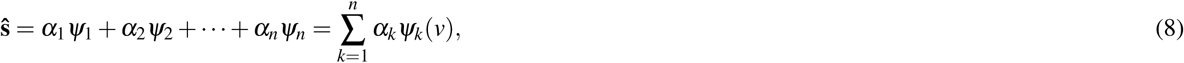

where the coefficient *α*_*k*_ of each functional harmonic *ψ*_*k*_ was estimated by projecting the task map **ŝ**(*v*) onto that particular harmonic *ψ*_*k*_. As such *α*_*k*_ are estimated as:

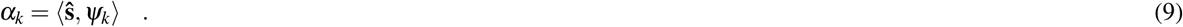

Then, each task map is reconstructed using Eq. 8. In this study, we limit our reconstructions to using a maximum of 100 non-constant functional harmonics (*n* = 101).

For a reconstruction **s**^*∗*^ _(*m*)_, where *m* indicates a binary vector of dimensionality 101 × 1 which contains ones for harmonic basis functions that are used in the reconstruction and zeros otherwise, we then compute the reconstruction error as:

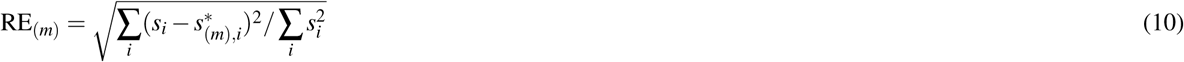

We also computed the Pearson correlations between **s** and **s**^*∗*^_(*m*)_. For comparing the correlations between task maps and reconstructions obtained from real functional harmonics versus randomized connectivity harmonics, we considered the number of comparisons to be nC = nTasks · nLevels, where the number of tasks equals 47 and the number of levels refers to the different numbers of harmonics used in the reconstructions, i.e. 0, 1, 2, 3, …, 20, 30, 40, …, 100, in 29 levels. From this we obtained a corrected alpha level of *α*_corr_ = 0.05*/*nC, and we computed the critical value as Fisher’s z-transform of the correlation which a sample has to exceed in order to be significantly higher than the random correlation:

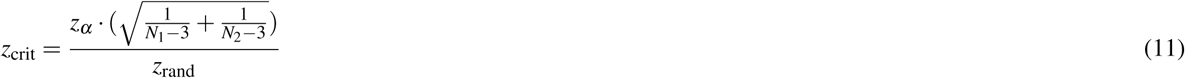

We obtain *z*_crit_ = 0.44, which corresponds to a minimum required empirical correlation of 0.41, with *N*_1_ = *N*_2_ = 59.412 (the number of vertices that contribute to the correlation values), *z*_*α*_ = 0.438 (the inverse Student’s t distribution with *N*_1_ = *N*_2_ = 59.412 degrees of freedom evaluated at 1 − *α*_corr_), and *z*_rand_ = atanh(0.05) (Fisher’s z-transform of the maximal random correlation between any reconstruction - with any number of functional harmonics - and any task).

### Visualization

#### Somatotopic areas

In the visual and somatosensory/motor cortices, functional harmonics are rather determined by retinotopy and somatotopy than by anatomical or microstructural features. For the former, somatotopic areas occupy exactly the same surface area as the sensorimotor core areas, 1, 2, 3a, 3b, and 4. We therefore replaced, where appropriate, the borders of the HCP parcellation by the borders of the five somatotopic regions.

#### Parcel borders for visualization

In order to discuss the meaning of the functional harmonics, we show borders of certain parcels on the cortical surfaces (Figure 2). We used three different methods to select which borders to show. First, for some functional harmonics, it was feasible to select these areas manually (for example, early visual areas in functional harmonic 4, somatotopic areas in functional harmonics 3 and 4). The anatomical supplementary information from Glasser et al. (2016)^3^ uses a functional grouping of many regions that we often used as a guideline, for instance to distinguish between early and association auditory cortex. Second, for some functional harmonics (for instance, functional harmonics 1 and 2), we show the borders of parcels that belong to resting state networks as defined by Yeo et al. (2011)^52^. The 7-network parcellation is provided by the HCP, which does not perfectly overlap with the HCP parcellation. We adjusted the network borders slightly to align the network borders to follow those of the parcels defined in HCP. Thereby we assigned each parcel to the RSN with which it had the most overlap. Third, some functional harmonics are too complex to manually select areas or networks (namely, functional harmonics 5, 6, 8, and 10). Here we employed simple k-means clustering on the functional harmonic, using k=2 (functional harmonics 5, 6, and 8) or k=3 (functional harmonic 10). To obtain meaningful clusters in the somatosensory/motor cortex, we again replaced the sensorimotor core regions 1, 2, 3a, 3b and 4 with the somatotopic areas. For this purpose, we used vertices within the core regions and re-assigned them to the somatotopic areas based on their distances to the sub-area borders.

## Author contributions statement

S. A. and K. G. designed the methodology and the analysis. J. P. and G. D. contributed to the design of the study. M. L. K. contributed to design of the methodology and the statistical analysis. M. L. K. and P. H. aided in the interpretation of the results. S. A., K. G. and M. L. K. wrote the manuscript. All authors reviewed the manuscript.

## Data availability

All data generated in this study are available from the corresponding author upon reasonable request.

## Code availability

All custom scripts used in this study are available from the corresponding author upon reasonable request.

## Additional information

**Competing financial interests** The authors declare no competing financial interests.

