## Supplementary Information for "Functional harmonics reveal multi-dimensional basis functions underlying cortical organization"

### Supplementary Material for
### Functional harmonics reveal multi-dimensional basis functions underlying cortical organization

**Katharina Glomb[1,*], Morten L. Kringelbach[2,3], Gustavo Deco[4,5,6,7], Patric Hagmann[1], Joel Pearson[8], and Selen Atasoy[2,3]**

[1]Department of Radiology, Lausanne University Hospital and University of Lausanne (CHUV-UNIL), Lausanne, Switzerland
[2]Department of Psychiatry, University of Oxford, Oxford, United Kingdom
[3]Center of Music in the Brain (MIB), Clinical Medicine, Aarhus University
[4]Center of Brain and Cognition, Universitat Pompeu Fabra, Barcelona, Spain
[5]ICREA, Institució Catalana de Recerca i Estudis Avancats (ICREA), Spain
[6]Department of Neuropsychology, Max Planck Institute for Human Cognitive and Brain Sciences, Leipzig, Germany
[7]School of Psychological Sciences, Monash University, Melbourne, Australia
[8]School of Psychology, University of New South Wales, Sydney Australia
[*]

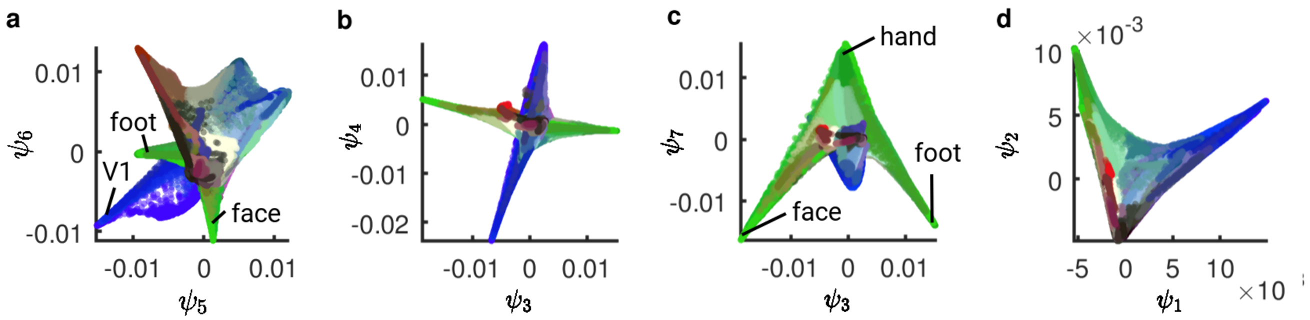


**Supplementary Figure 1.** Two-dimensional subspaces formed by pairs of functional harmonics, referred to as $\psi$. The color code is taken from the HCP parcellation[1], with blue corresponding to visual areas, red, auditory areas, and green, somatosensory/motor areas, and the color gradient running from white to black signifying task-positive to task-negative areas. **a**: Two-dimensional subspace formed by functional harmonics 5 and 6 ($\psi_5$ and $\psi_6$). V1 (blue), two somatotopic areas (hand and face, green) and auditory areas (red) are separated from each other and from remaining areas. **b**: Two-dimensional subspace formed by functional harmonics 3 and 4 ($\psi_3$ and $\psi_4$). Visual (blue) and somatosensory/motor/auditory (green and red) systems are orthogonal to each other. **c**: Two-dimensional subspace formed by functional harmonics 3 and 7 ($\psi_3$ and $\psi_7$). A different separation of somatotopic regions from that shown in Figure 3a is apparent. **d**: Two-dimensional subspace formed by functional harmonics 1 and 2 ($\psi_1$ and $\psi_2$). This reproduces findings from[2], where visual areas (blue), somatosensory/motor areas (green) and higher-order areas belonging to the default mode network (black) are separated from each other.

Figure 2, top, shows the "somatotopy index", which quantifies to which degree somatotopic areas (face, eye, hand, trunk, foot) are well-circumscribed by the functional harmonics. We use the Euclidean distances on the functional harmonics (see Online Methods for details). We plot this "somatotopy index" for each somatotopic region on the original functional harmonics (circles), compared to spherical rotations (gray crosses, 300 rotations). Higher indices mean a clearer separation, and the results correspond to what can be visually appreciated in Figure 2. For example, the separation of the hand-areas is especially strong in functional harmonic 11.

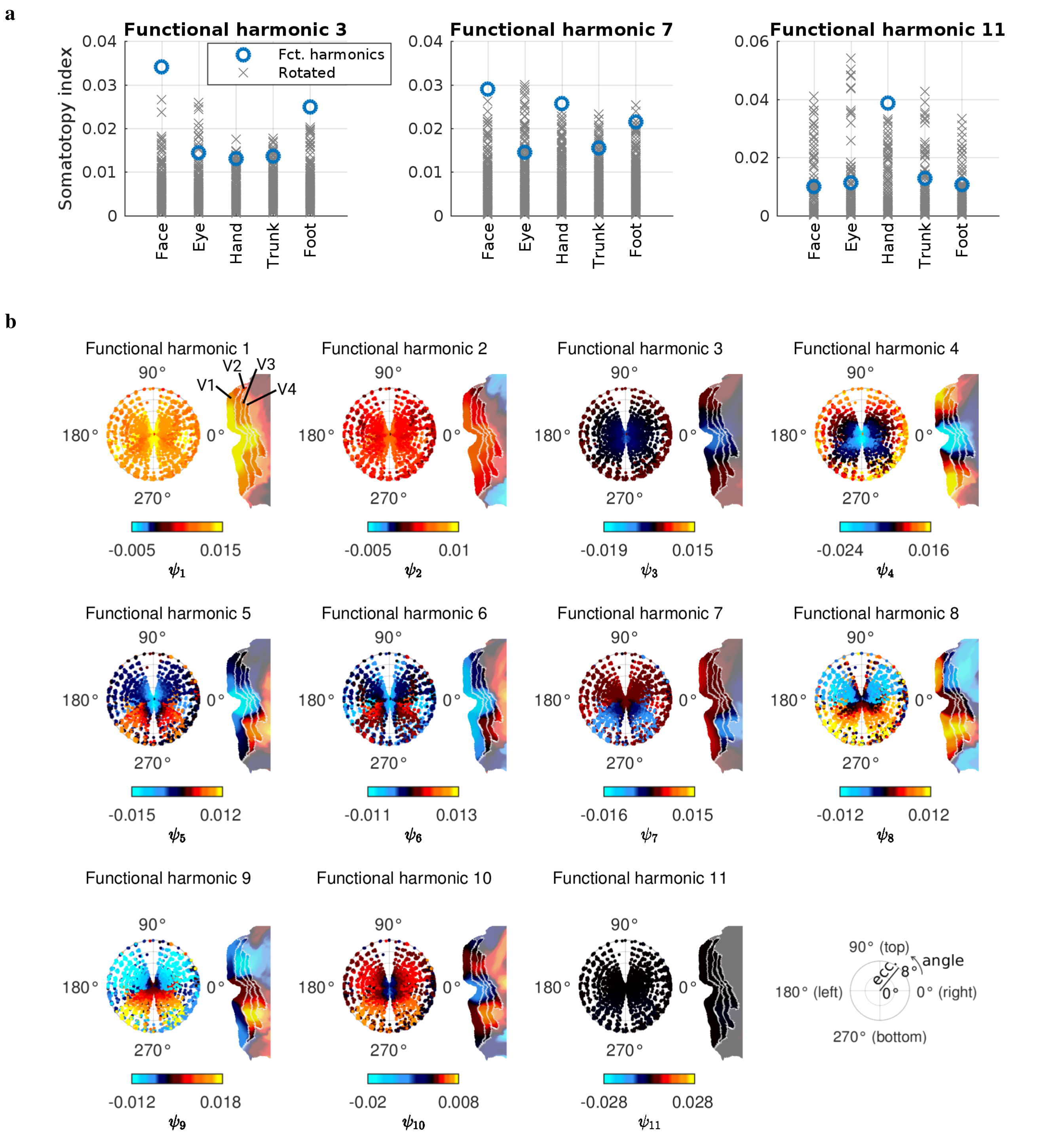


**Supplementary Figure 2. a**: “Somatotopy index" for functional harmonics 3, 7, and 11, and each of the somatotopic subregions (averaged across hemispheres), blue circles; grey crosses are computed from 300 sets of rotations of functional harmonics. This index will be high if the region is separated well from both the entire rest of the cortex and the other somatotopic regions (see Methods). **b**: All retinotopic mappings in V1-V4 found in the first 11 harmonics ($\psi_1$-$\psi_{11}$).

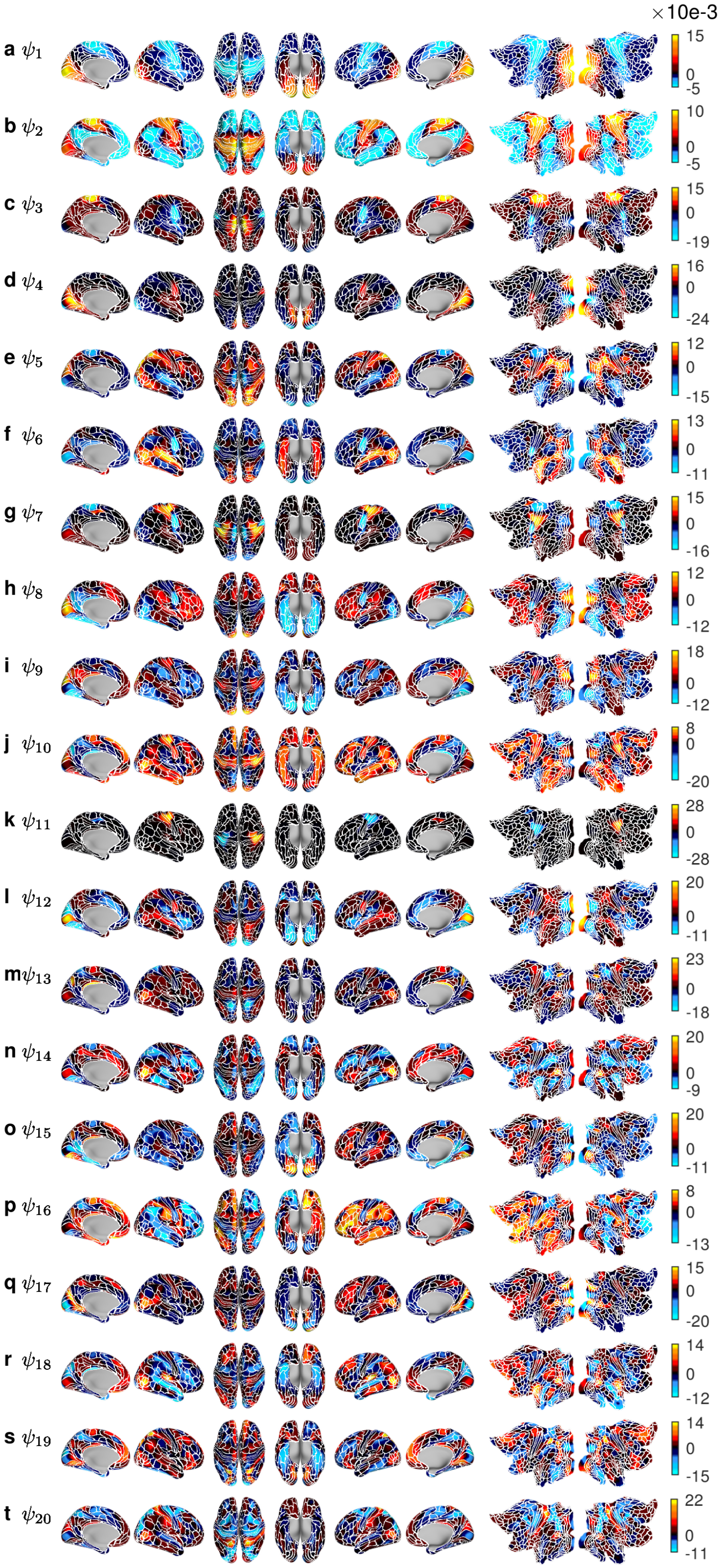


**Supplementary Figure 3.** First 20 functional harmonics (referred to as $\psi_1$-$\psi_{20}$) with all borders defined in the HCP parcellation[1].

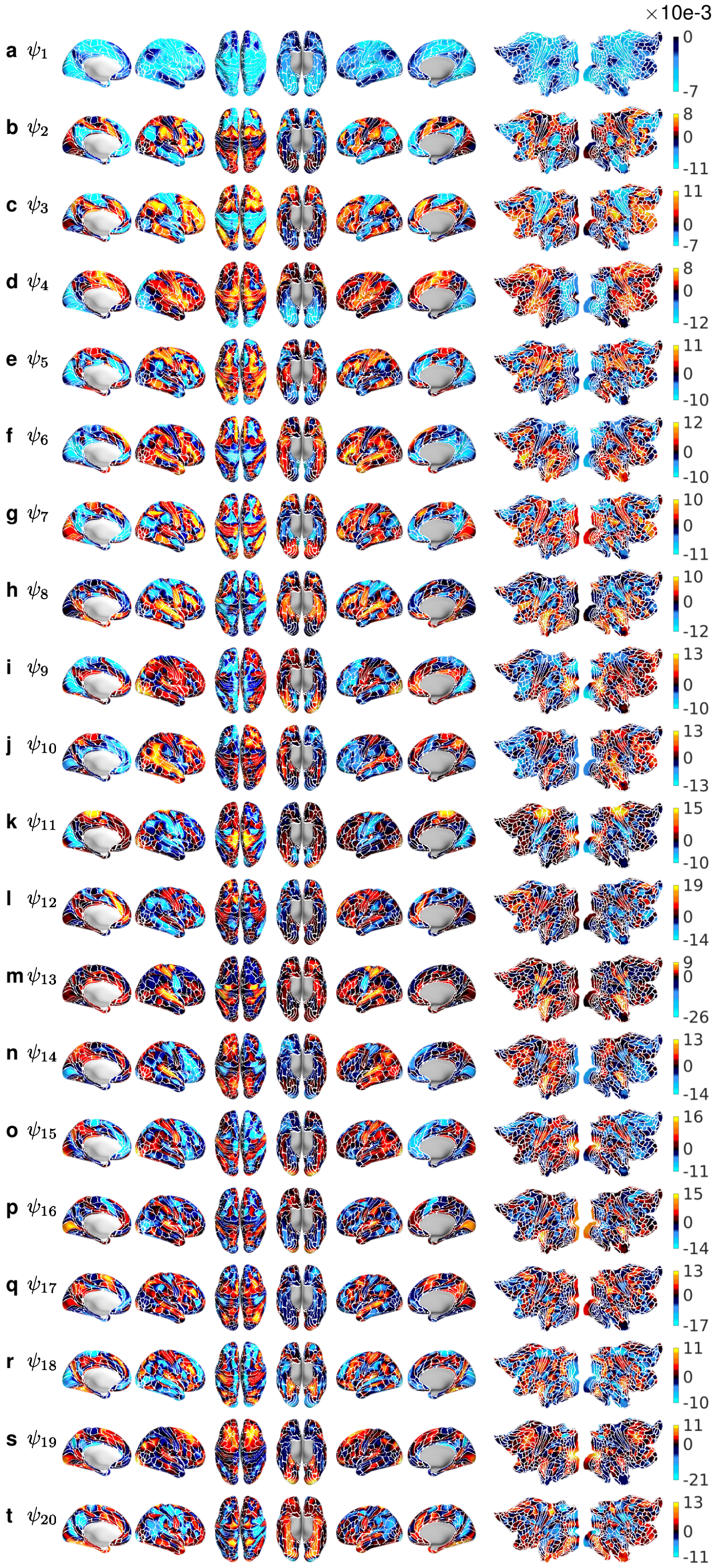


**Supplementary Figure 4.** First 20 eigenvectors derived from the dense functional connectivity matrix (referred to as $\psi_1$-$\psi_{20}$) with all borders defined in the HCP parcellation[1].

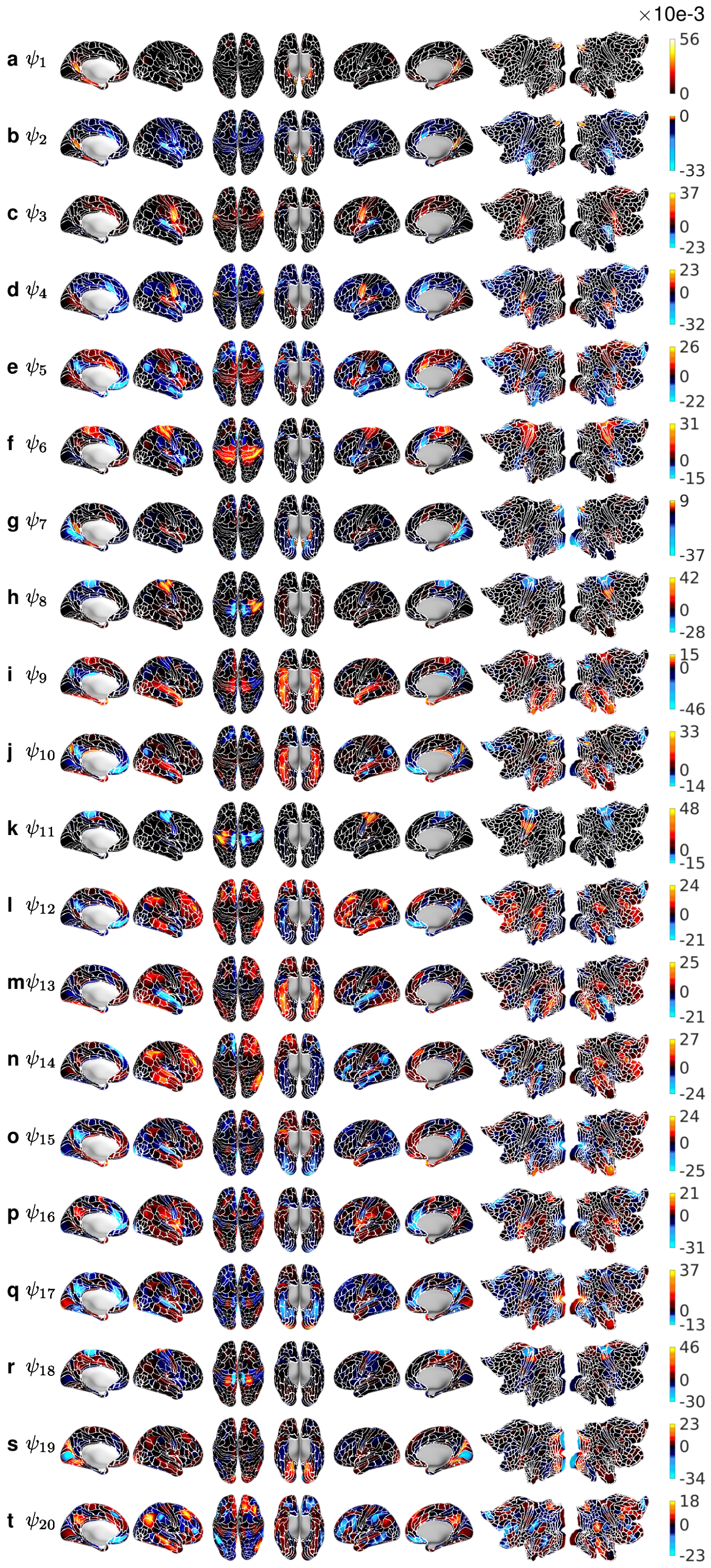


**Supplementary Figure 5.** First 20 eigenvectors derived from the adjacency matrix (referred to as $\psi_1$-$\psi_{20}$) with all borders defined in the HCP parcellation[1].

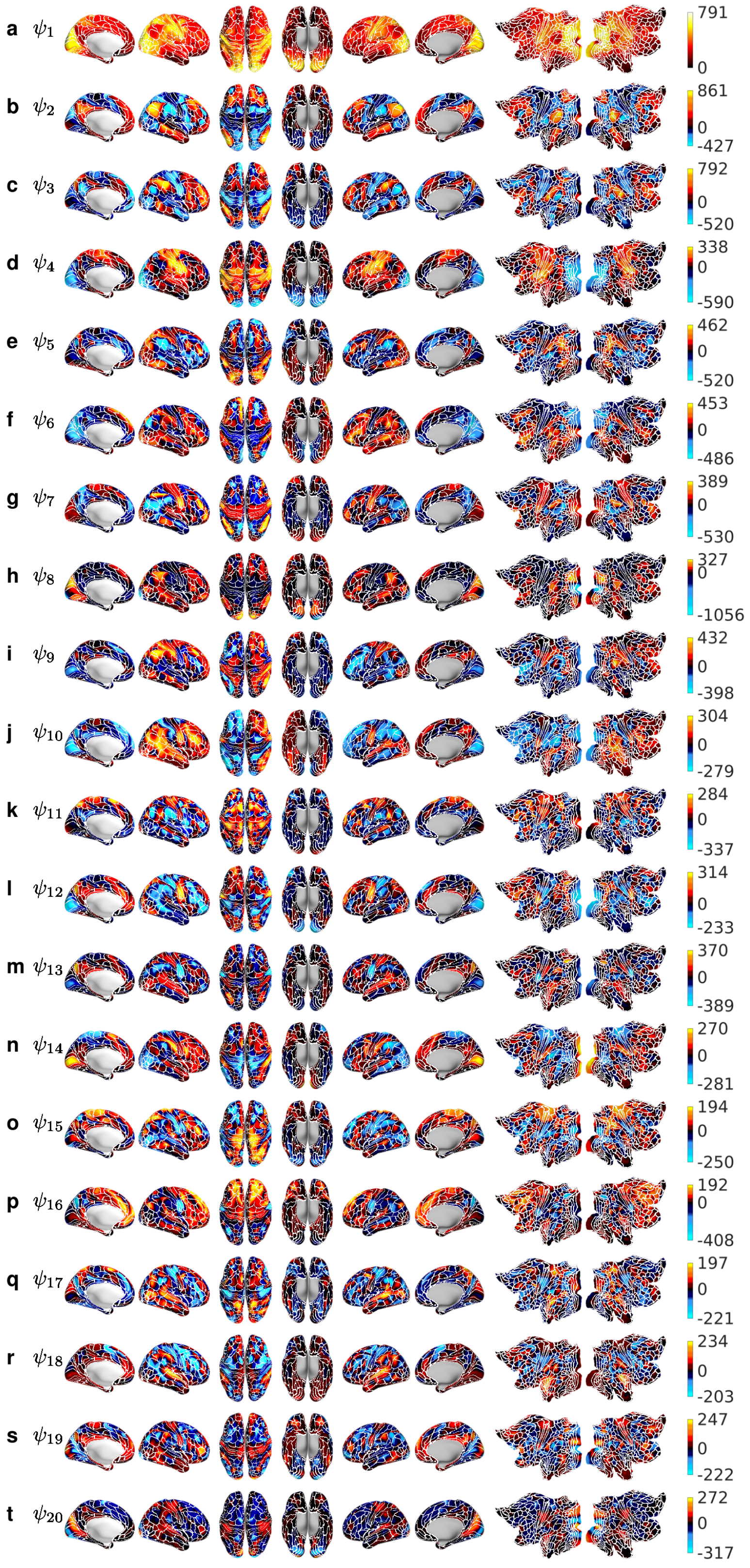


**Supplementary Figure 6.** First 20 principal components, i.e. eigenvectors of the covariance matrix, as provided by the HCP (referred to as $\psi_1$-$\psi_{20}$) with all borders defined in the HCP parcellation[1].

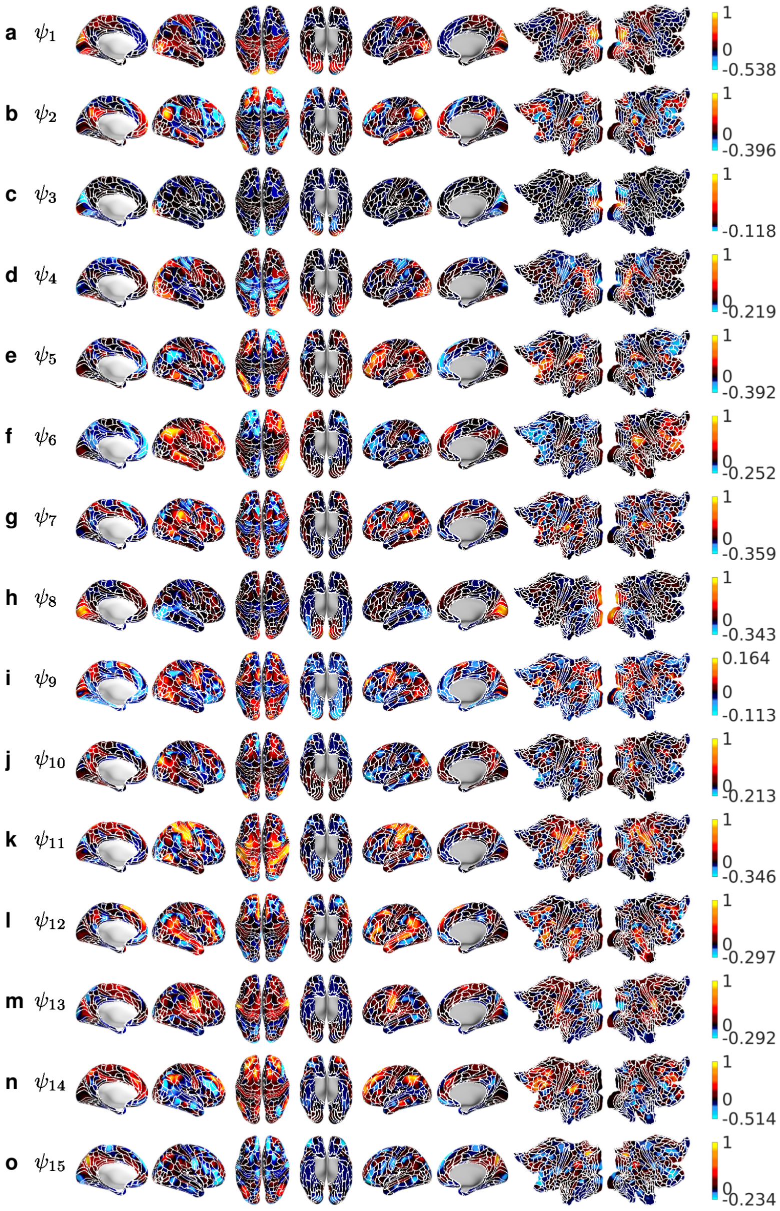


**Supplementary Figure 7.** 15 independent components as provided by the HCP (referred to as $\psi_1$-$\psi_{15}$) with all borders defined in the HCP parcellation[1]. In this case, the number of components was set to 15, i.e. the figure shows the complete set of ICs.

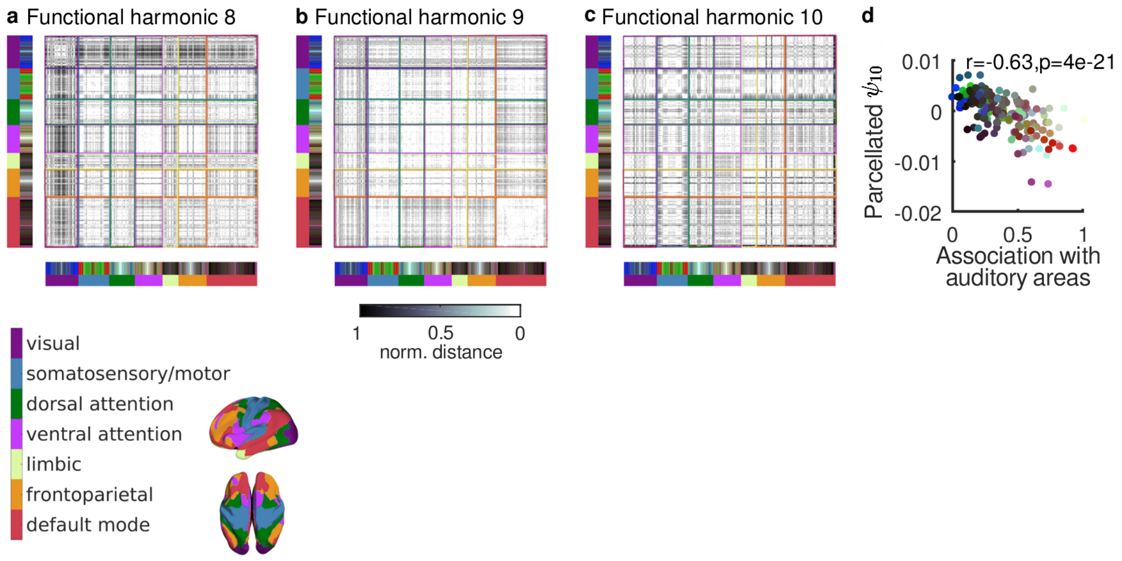


**Supplementary Figure 8.** Normalized average Euclidean distances between all pairs of parcels, ordered by resting state network (RSN) membership according the Yeo 7-Network parcellation[3] (see legend), for functional harmonics 8 (panel **a**), 9 (panel **b**), and 10 (panel **c**). **d** Correlation between the degree to which areas are related to auditory regions[1] and the value of functional harmonic 10 ($\psi_{10}$), averaged within each of the 360 parcels. The color code is taken from the parcellation in Glasser et al. (2016)[1], see also Figure 1**d**.

Figure 8a-c illustrates how functional harmonics 8-10 exhibit different divisions between higher order networks defined by the 7-network resting state network (RSN) parcellation by Yeo and colleagues[3]. In functional harmonic 8 (Figure 8a), the distances within the ventral attention network (vATT) are very small, as well as distances between regions belonging to the vATT and the frontal parietal network (FPN). As shown in Figure 4d, we observe a strong retinotopic gradient across regions V1-V4, which is reflected by high distances within the visual RSN.

In functional harmonic 9 (Figure 8b), most RSNs show small within-network distances, particularly the somatosensory/motor, dorsal and ventral attention, as well as default mode networks. Small distances also occur between the somatosensory/motor and default mode networks.

In functional harmonic 10 (Figure 8c), only the ventral attention and limbic networks exhibit small within- (but not between-) network distances. We also observed that functional harmonic 10 ($\psi_{10}$) captures the hierarchical organization of the auditory system (Figure 8d). To quantify this agreement, we measured the correlation between the spatial pattern of functional harmonic 10 ($\psi_{10}$) and the extent to which each area is associated with the auditory network in the resting state (degree of auditory involvement)[1]. We found a significant correlation ($r = -0.63$, $p = 4 \cdot 10^{-21}$) between functional harmonic 10 ($\psi_{10}$) and the degree of auditory involvement of the functional areas (Figure 3b). Also, certain regions of the frontoparietal network appear to be strongly separated from all other networks (black bands within frontoparietal network running across all other networks).

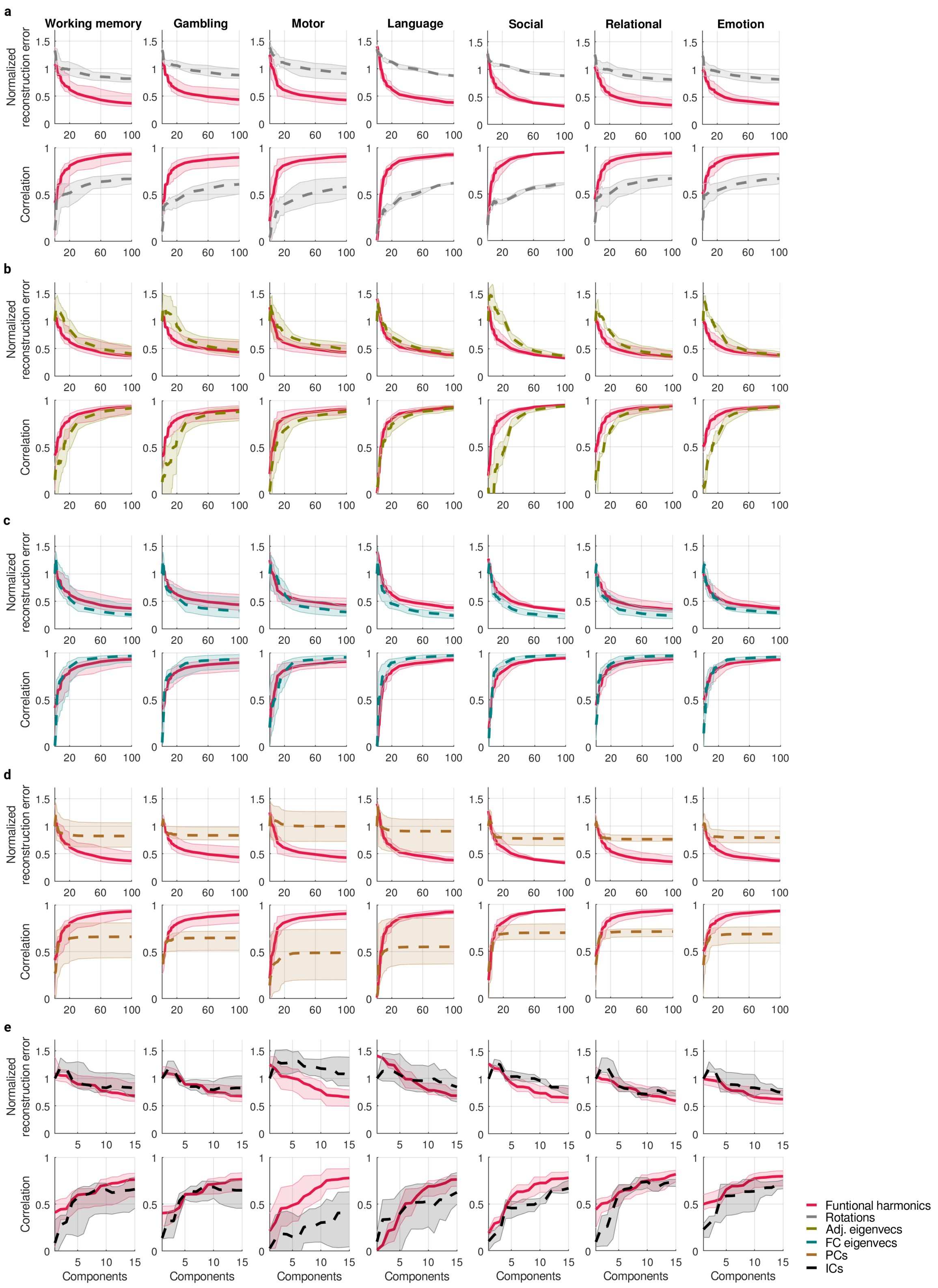


**Supplementary Figure 9.** Reconstruction performance of functional harmonics (solid lines) compared to **a**: their rotations, **b**: eigenvectors of the adjacency matrix, **c**: eigenvectors of the dense FC matrix, **d**: principal components (PCs), **e**: independent components (ICs; dashed lines). Shaded areas show the range (minimum and maximum) of reconstructions errors and (first row of each panel) correlations (second row of each panel) across all tasks in this group.

Figure 10 shows, for the 7 task groups used by the HCP[4], all task maps (left columns) with the normalized power spectra over the first 11 non-constant functional harmonics (middle columns), and the reconstructed maps using the strongest, four strongest, and forty strongest contributing functional harmonics (right columns). In cases where the strongest contributing functional harmonic was the "0th", i.e. the constant functional harmonic, we show the 2nd-strongest, i.e. the strongest non-constant functional harmonic in the right column. This is case in Figure 10e, v, w, x, z, da, fa, ja, ka, ma, and na. In those cases, the maximum value in the polar plot in the middle column may not be 1, because the "0th" functional harmonic is not shown in these plots. Note that however the reconstructions using more than one functional harmonic do use the constant functional harmonic. Importantly, there was no task whose strongest contributing functional harmonic was beyond the 11th non-constant functional harmonic.

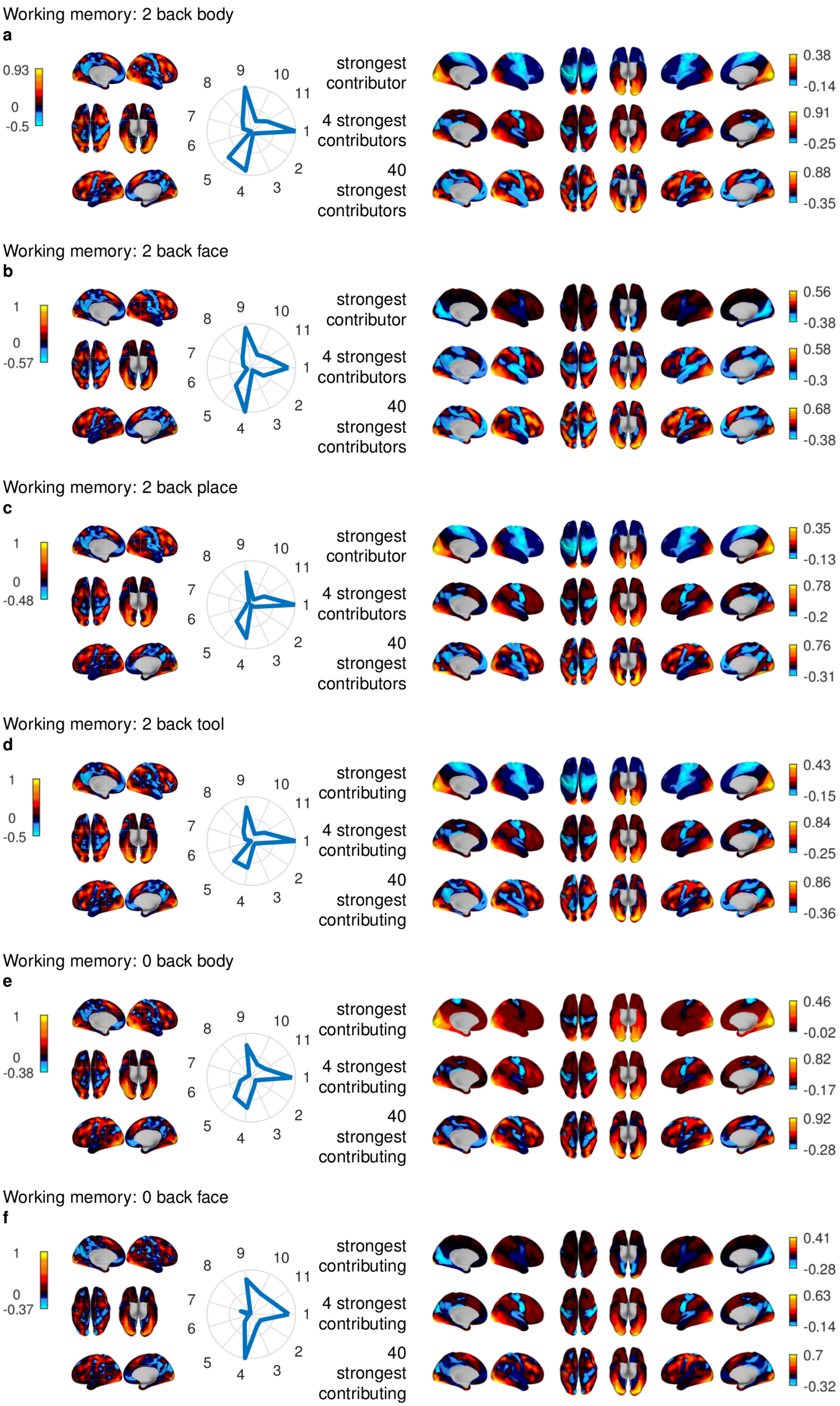


**Supplementary Figure 10**

Working memory: 0 back place

**g**

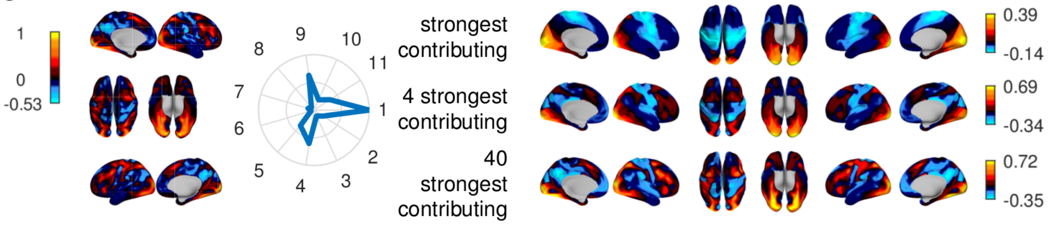


Working memory: 0 back tool

**h**

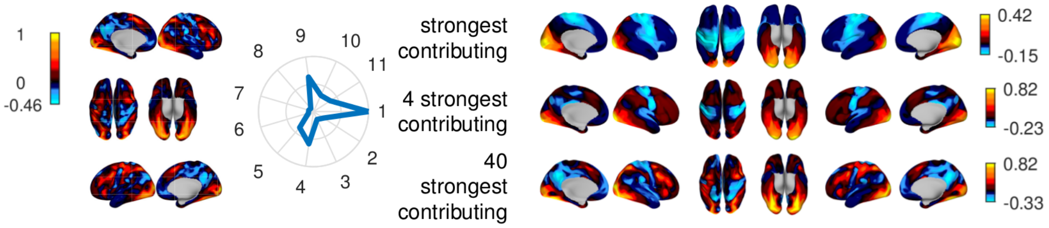


Working memory: average 2 back

**i**

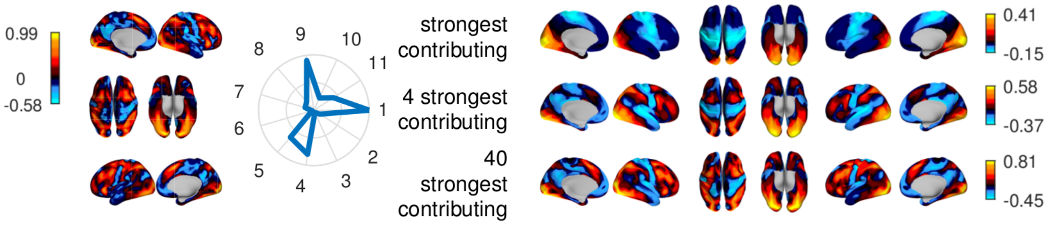


Working memory: average 0 back

**j**

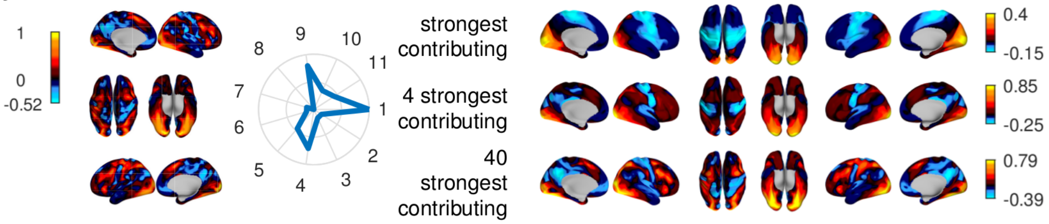


Working memory: contrast 2 back minus 0 back

**k**

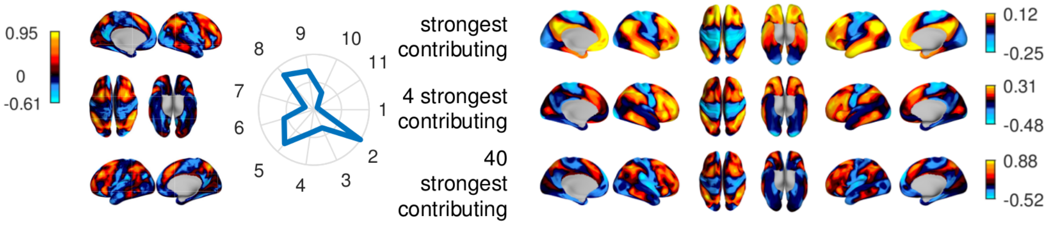


Working memory: average body

**l**

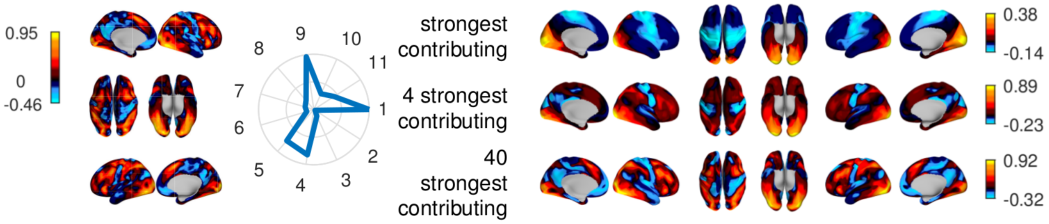


**Supplementary Figure 10.** (continued)

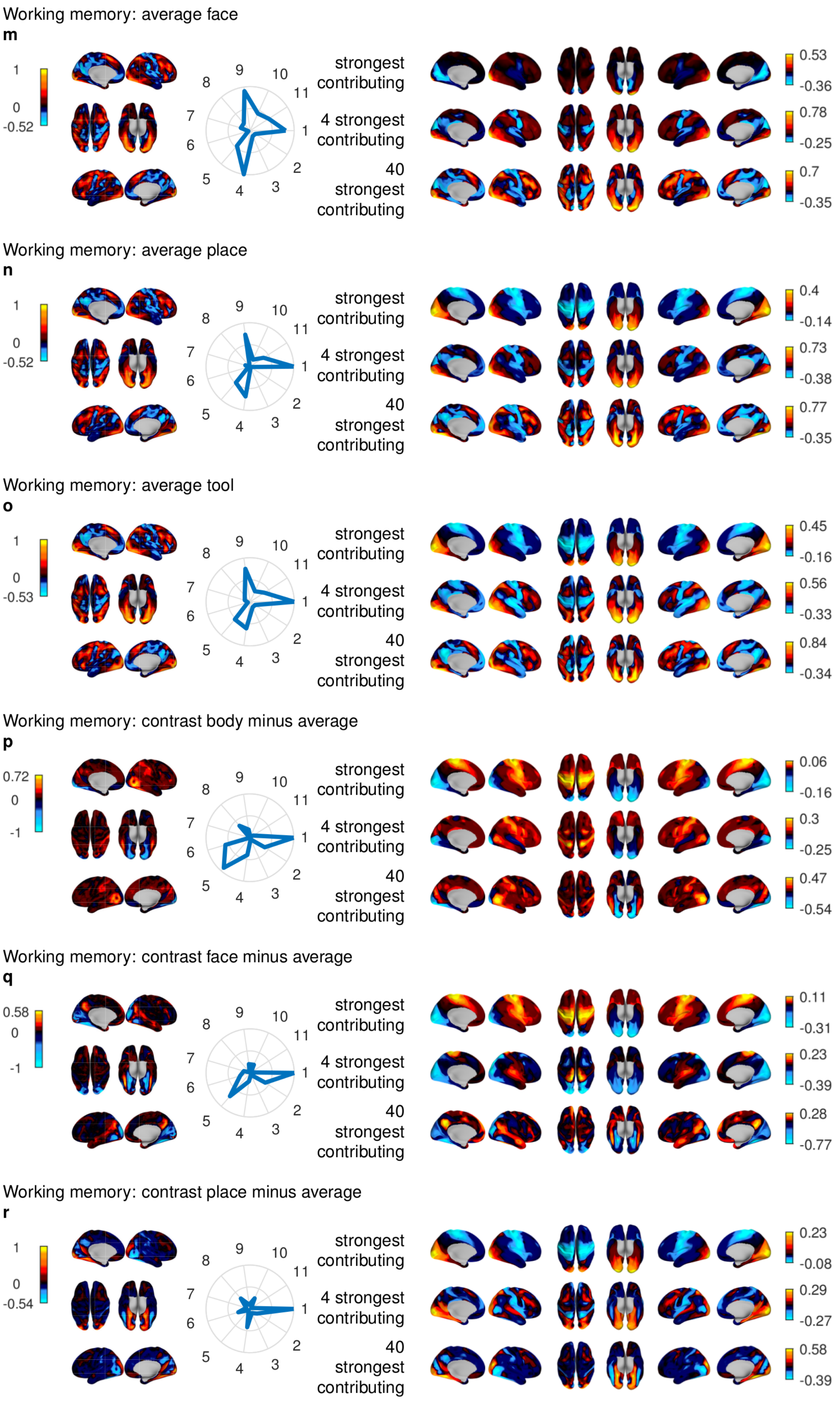


**Supplementary Figure 10.** (continued)

Working memory: contrast tool minus average

**s**

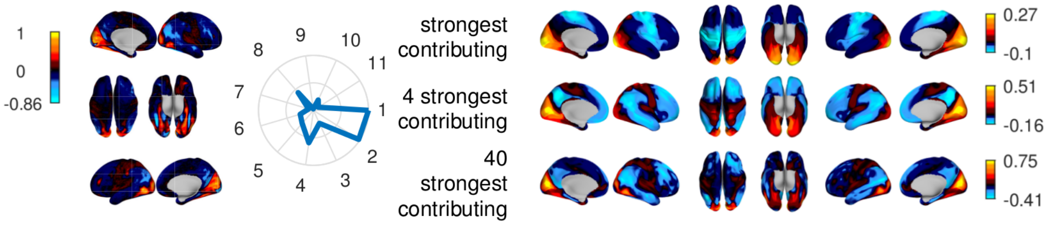


Gambling: punish

**t**

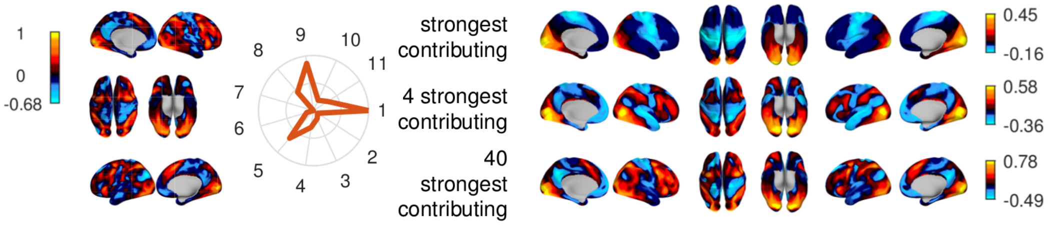


Gambling: reward

**u**

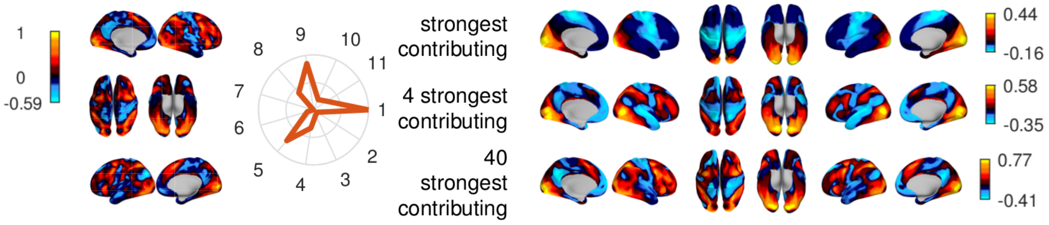


Gambling: contrast punish minus reward

**v**

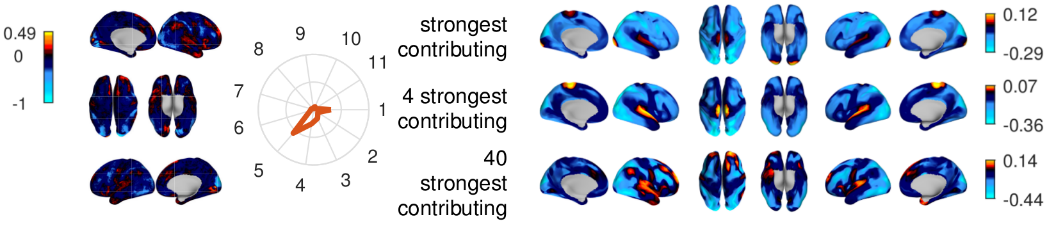


Motor: cue

**w**

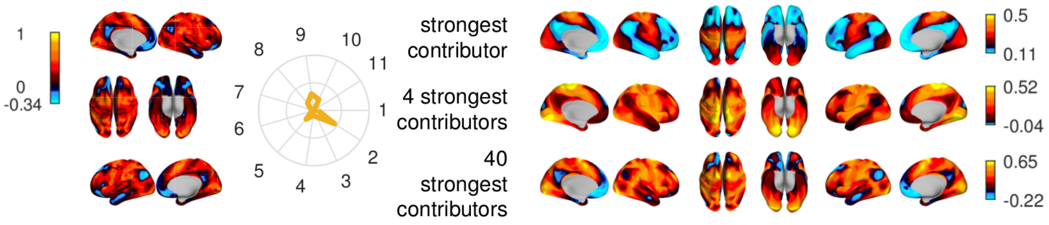


Motor: left foot

**x**

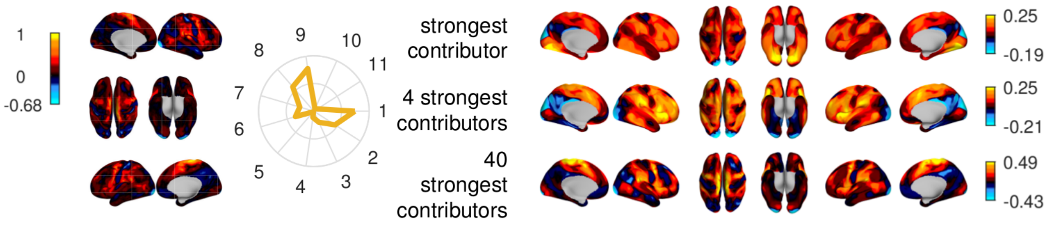


**Supplementary Figure 10.** (continued)

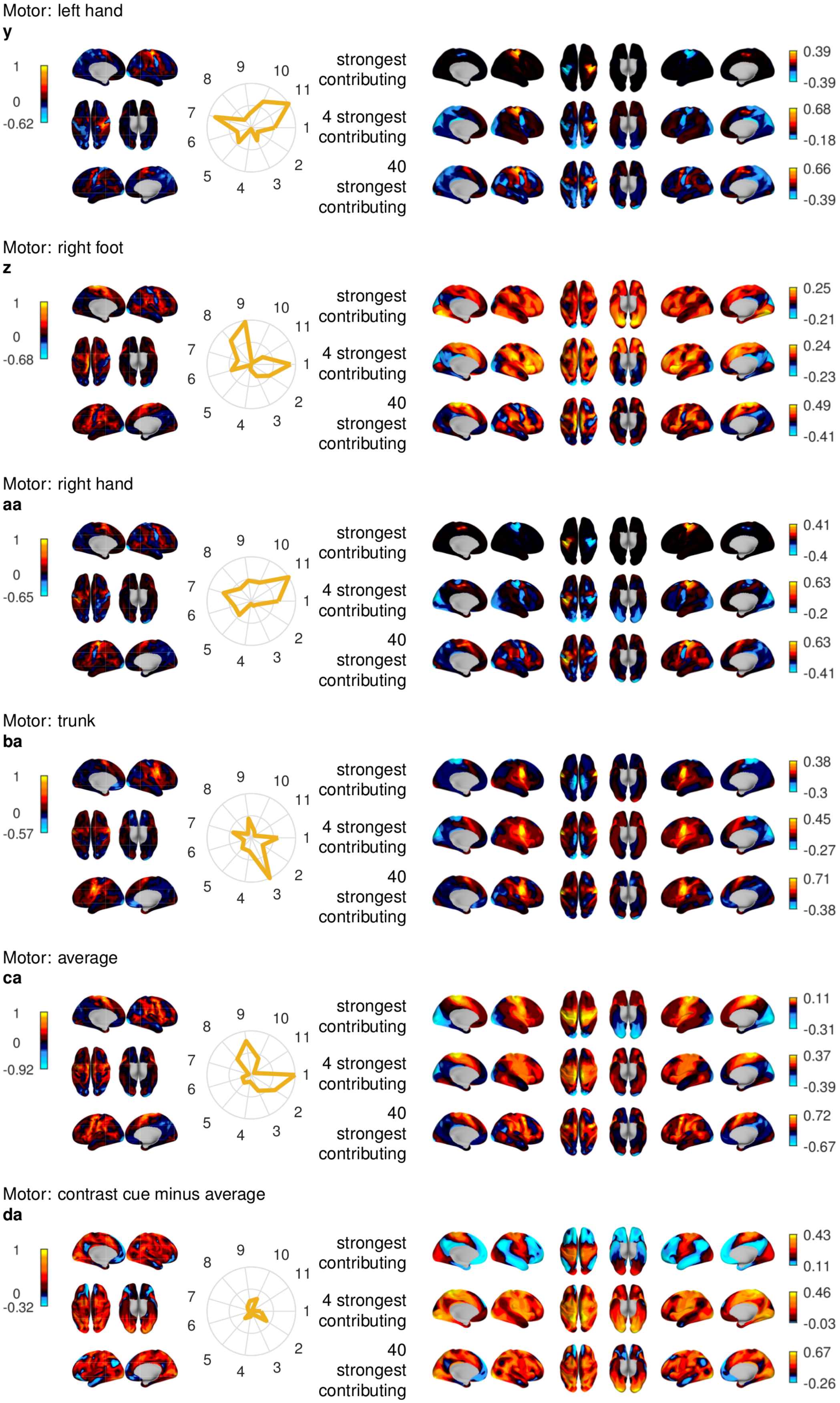


**Supplementary Figure 10.** (continued)

Motor: contrast left foot minus average
**ea**
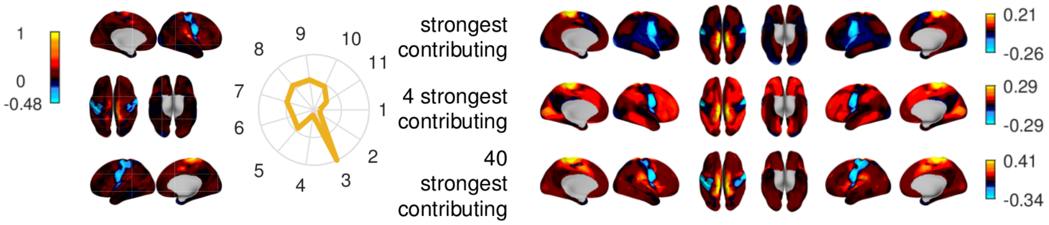


Motor: contrast left hand minus average
**fa**
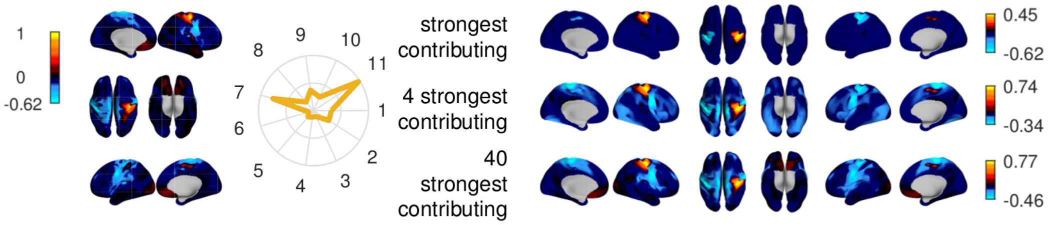


Motor: contrast right foot minus average
**ga**
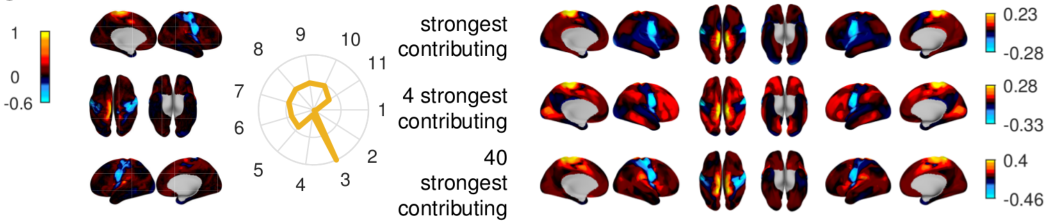


Motor: contrast right hand minus average
**ha**
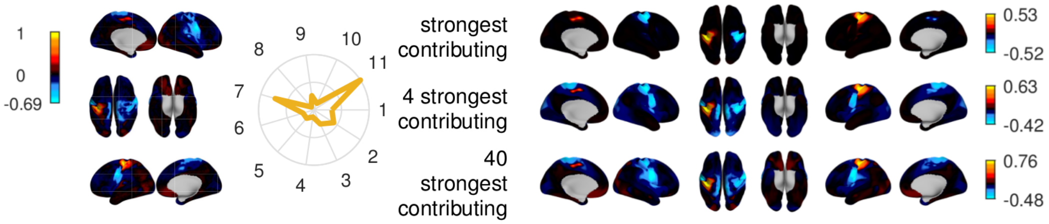


Motor: contrast trunk minus average
**ia**
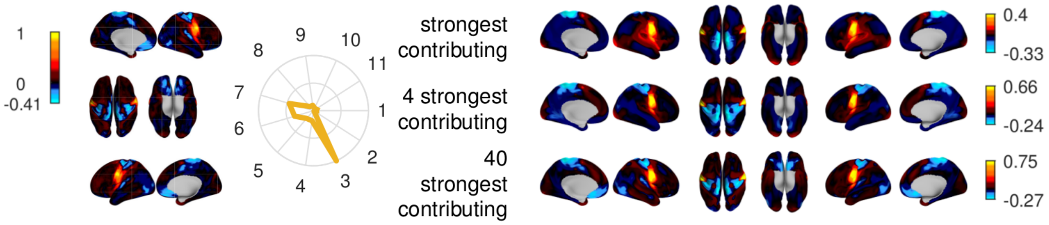


Language: math
**ja**
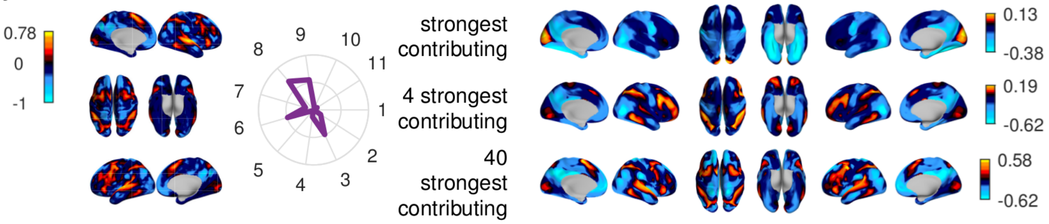


**Supplementary Figure 10.** (continued)

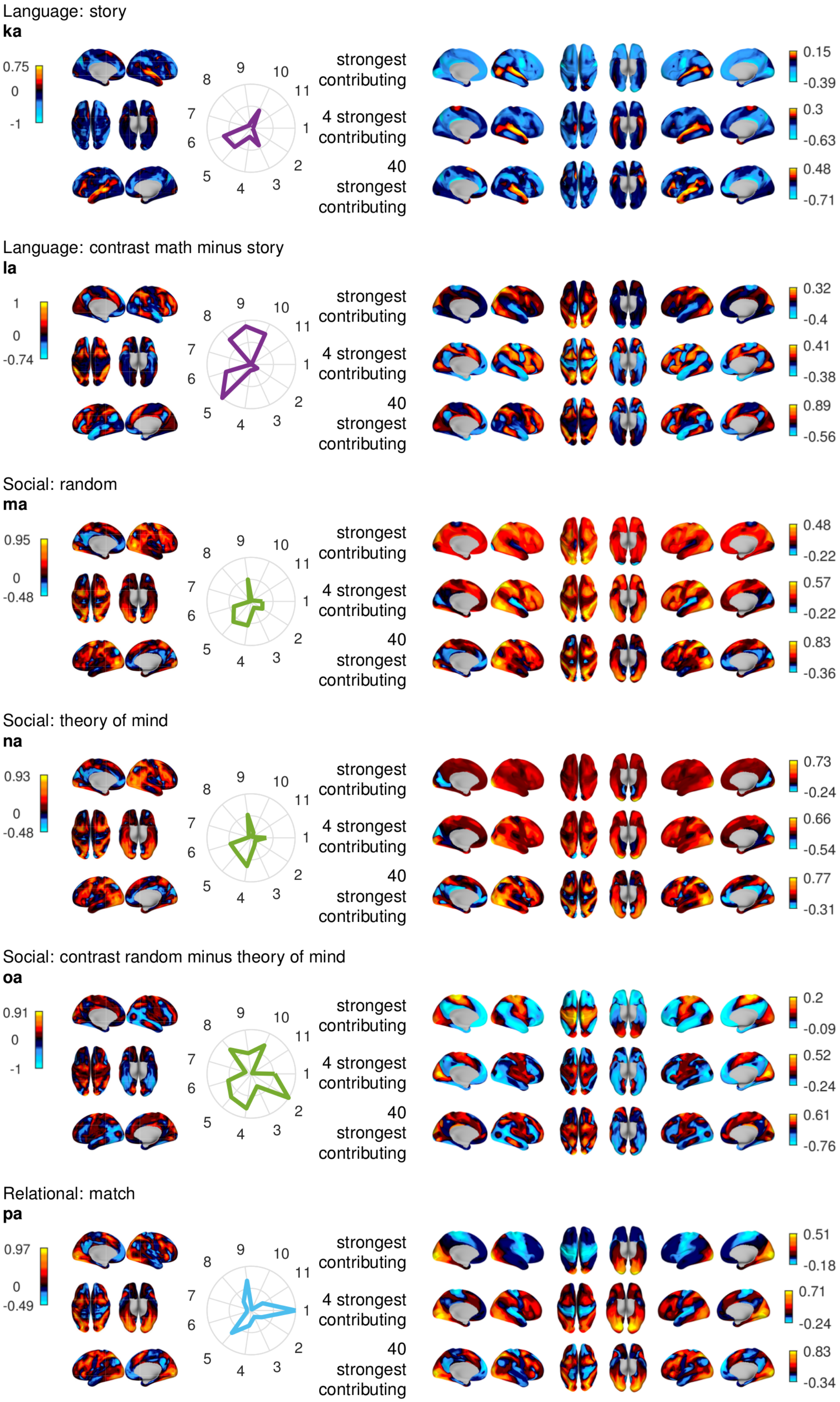


**Supplementary Figure 10.** (continued)

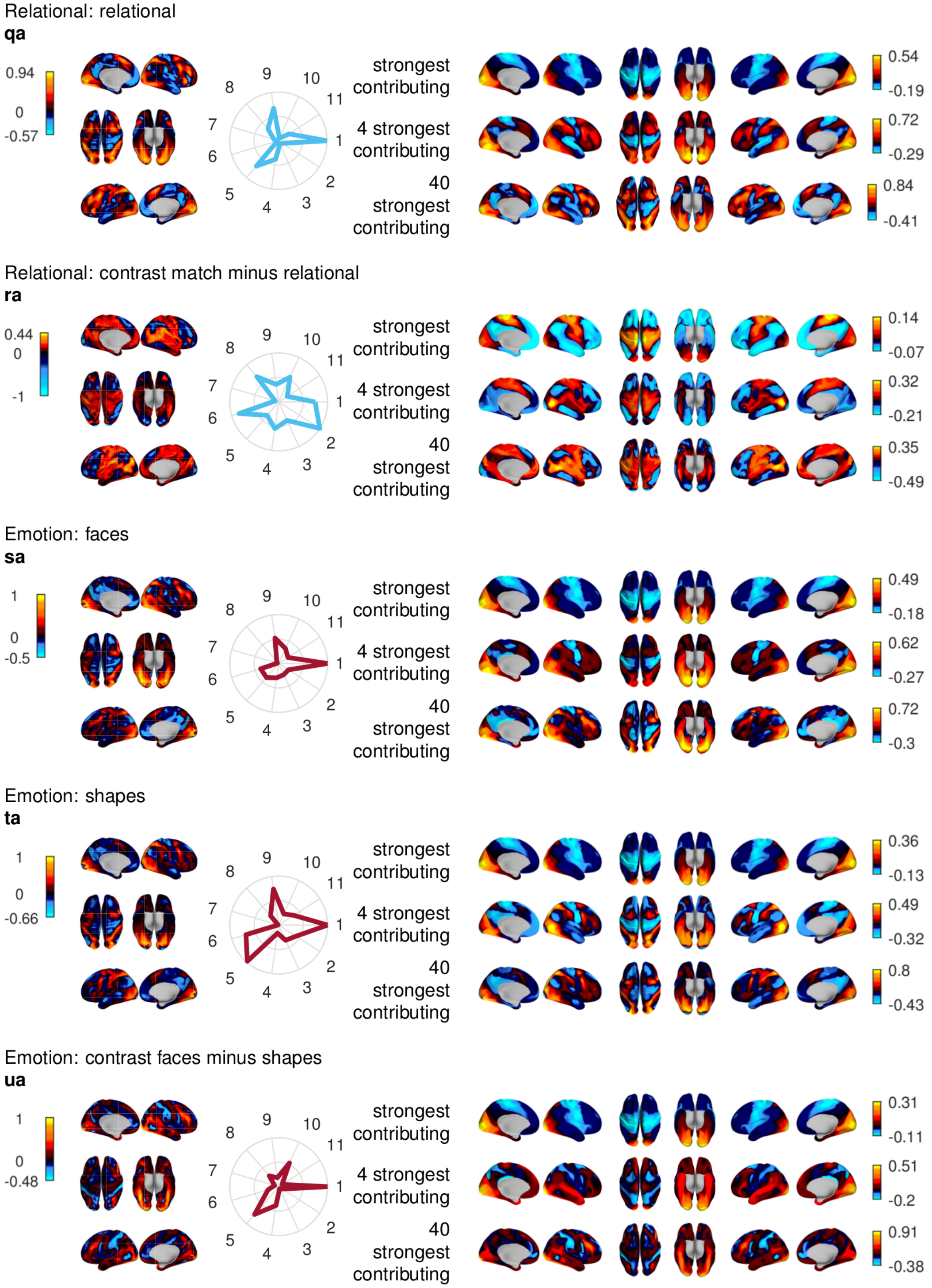


**Supplementary Figure 10.** (continued) Reconstructions of task maps. The left column shows the original task map provided by the HCP[4], see table 1 in Online Methods; the middle column, the normalized (over all 101 functional harmonics that were computed, i.e. the constant one and the first 100 non-constant ones) absolute value of the coefficient which measures the contribution of the first non-constant harmonics; the right column shows the reconstructions of the task maps using the indicated number of strongest contributing functional harmonics. **a-s**: Working memory, **t-v**: Gambling, **w-ia**: Motor, **ja-la**: Language, **ma-oa**: Social, **pa-ra**: Relational, **sa-ua**: Emotion.
